# scBSP: A fast and accurate tool for identifying spatially variable genes from spatial transcriptomic data

**DOI:** 10.1101/2024.05.06.592851

**Authors:** Jinpu Li, Yiqing Wang, Mauminah Azam Raina, Chunhui Xu, Li Su, Qi Guo, Qin Ma, Juexin Wang, Dong Xu

## Abstract

Spatially resolved transcriptomics have enabled the inference of gene expression patterns within two and three-dimensional space, while introducing computational challenges due to growing spatial resolutions and sparse expressions. Here, we introduce scBSP, an open-source, versatile, and user-friendly package designed for identifying spatially variable genes in large-scale spatial transcriptomics. scBSP implements sparse matrix operation to significantly increase the computational efficiency in both computational time and memory usage, processing the high-definition spatial transcriptomics data for 19,950 genes on 181,367 spots within 10 seconds. Applied to diverse sequencing data and simulations, scBSP efficiently identifies spatially variable genes, demonstrating fast computational speed and consistency across various sequencing techniques and spatial resolutions for both two and three-dimensional data with up to millions of cells. On a sample with hundreds of thousands of sports, scBSP identifies SVGs accurately in seconds to on a typical desktop computer.

## INTRODUCTION

Spatially Resolved Transcriptomics (SRT) has evolved rapidly in biological and biomedical research (1–4). Early technologies in SRT, such as single-molecule fluorescence in situ hybridization (smFISH) (5) and sequencing-based approaches (e.g., 10X Visium), were primarily designed for small samples on two-dimensional (2D) slices with relatively low spatial resolution. In contrast, recent advancements, notably in three-dimensional (3D) techniques like STARmap (6), have demonstrated substantial advantages in a more comprehensive and faithful representation of intact organ structures and functions, enhancing accurate quantitative interpretation (6–12). Furthermore, advancements in high-resolution transcriptomics, including Slide-seq (13), high-definition spatial transcriptomics (HDST) (14), 10X Xenium, NanoString CosMx, MERSCOPE, Stereo-seq (15) and other sequencing approaches have propelled transcriptome-wide profiling to a single-cell or subcellular resolution with a greater number of spots or cells (16, 17). This progress provides a more accurate and comprehensive measurement of tissue structures, marking a significant stride forward in SRT capabilities (18, 19). However, the shift from 2D to 3D space and substantial increase in spatial resolution brings new computationally challenges in data analysis, including the significantly growing number of spatial locations and a high proportion of zero values, limited sequencing depth (20–23).

A common SRT analysis is to identify Spatially Variable Genes (SVGs), which play a crucial role in comprehending tissue structures and functions (24–30). Unlike the analysis of differentially expressed genes on traditional RNA-seq, identifying SVGs in SRT involves considering spatial context and geometric coordinates. Numerous methods have been developed for the SVG detection in 2D space since 2018 (21, 31–38). Beyond SVG detection in 2D samples, preliminary research has been conducted on 3D SRT, including SPARK-X (21) and spaGCN (36). However, substantial challenges persist in identifying SVGs in large-scale SRT as the number of spots reaches millions.

Methods like SpatialDE (37) and SPARK (38) were specifically developed for detecting SVGs in small samples. To enhance computational efficiency, two hybrid methods, nnSVG (34) and SOMDE (35), were optimized to scale linearly with the increasing number of spatial locations by incorporating graph and kernel approaches, but their computational efficiency is still insufficient to deal with SRT data featuring cellular or sub-cellular resolution, typically encompassing tens or hundreds of thousands of spatial locations with thousands of genes. Additionally, the violation of model assumptions and computational complexity hinders the application of current methods. For example, due to inconsistencies in inter-plane and within-plane spatial resolution, identifying SVGs from 3D SRT has limited accuracies.

Our recent method, BSP (39), has introduced a dimension-agnostic and granularity-based solution by calculating the statistical discrepancies between a paired of big and small patch. Its model performance in identifying SVGs was validated in both simulations and experimental studies utilizing 2D and 3D data. This approach showcased reasonable computational efficiency, taking only 7 and 18 minutes on a typical Ubuntu 6.04.4 LTS workstation with Intel(R) Xeon(R) W-2125 CPU (4.00 GHz and 32 GB memory) for the Slide-seq and Slide-seq V2 data on the mouse cerebellum, comprising 18,671 genes on 25,551 beads and 23,096 genes on 39,496 beads, respectively. However, when confronted with data featuring higher spatial resolution, such as the HDST olfactory bulb data that encompassing 19,950 genes measured on 181,367 spots, the original BSP algorithm requires 4 hours and 90GB of memory on a High-Performance Computer equipped with a 2.20 GHz Intel Xeon(R) CPU E5-2699 v4. This increased computational time and memory usage impose limitations on the applicability of BSP in 3D and large-scale SRT.

Here we present scBSP (single-cell big-small patch), a significantly enhanced version of BSP, to address computational challenges in the identification of SVGs from large-scale two/three-dimensional SRT data. scBSP selects a set of neighboring spots within a certain distance to capture the regional means and filters the SVGs using the velocity of changes in the variances of local means with different granularities. Incorporating sparse matrix techniques and approximate nearest neighbor search, scBSP significantly reduces computational time and memory consumption compared to the original implementation of

BSP and other existing approaches, reducing the processing time for a high-definition spatial transcriptomics data for 19,950 genes measured on 181,367 spots with 5.95 seconds. Applied to diverse sequencing data and simulations of thousands of genes from tens or hundreds of thousands of spots, scBSP accurately identifies SVGs within seconds to minutes on a typical laptop. Notably, scBSP stands out for its robustness, as it does not assume any specific distribution of gene expression levels or spatial patterns of spots and it does not have model parameters to tune or train. This feature ensures its adaptability to various SRT sequencing techniques, ensuring consistent results without the need for adapting the model and effectiveness in addressing the zero-inflation issue commonly encountered in large-scale data analyses.

## MATERIAL AND METHODS

### Overview of scBSP workflow

scBSP is an open-source R package and a corresponding Python library that implements a spatial granularity-based algorithm for identifying SVGs in dimension-agnostic and technology-agnostic SRT data. A pair of patches is defined for each spot in the SRT data, one of which comprise neighboring spots within a small radius and the other within a large radius. The average expression within each patch is calculated, reflecting the local expression levels at various spatial granularities. The variance of local expressions is computed across all spots for each given radius. The key concept of this approach is to capture the rate of change in variance of local means as the granularity level increases, measured by the variance ratio with different patch sizes (Figure 1).

**Figure 1:**
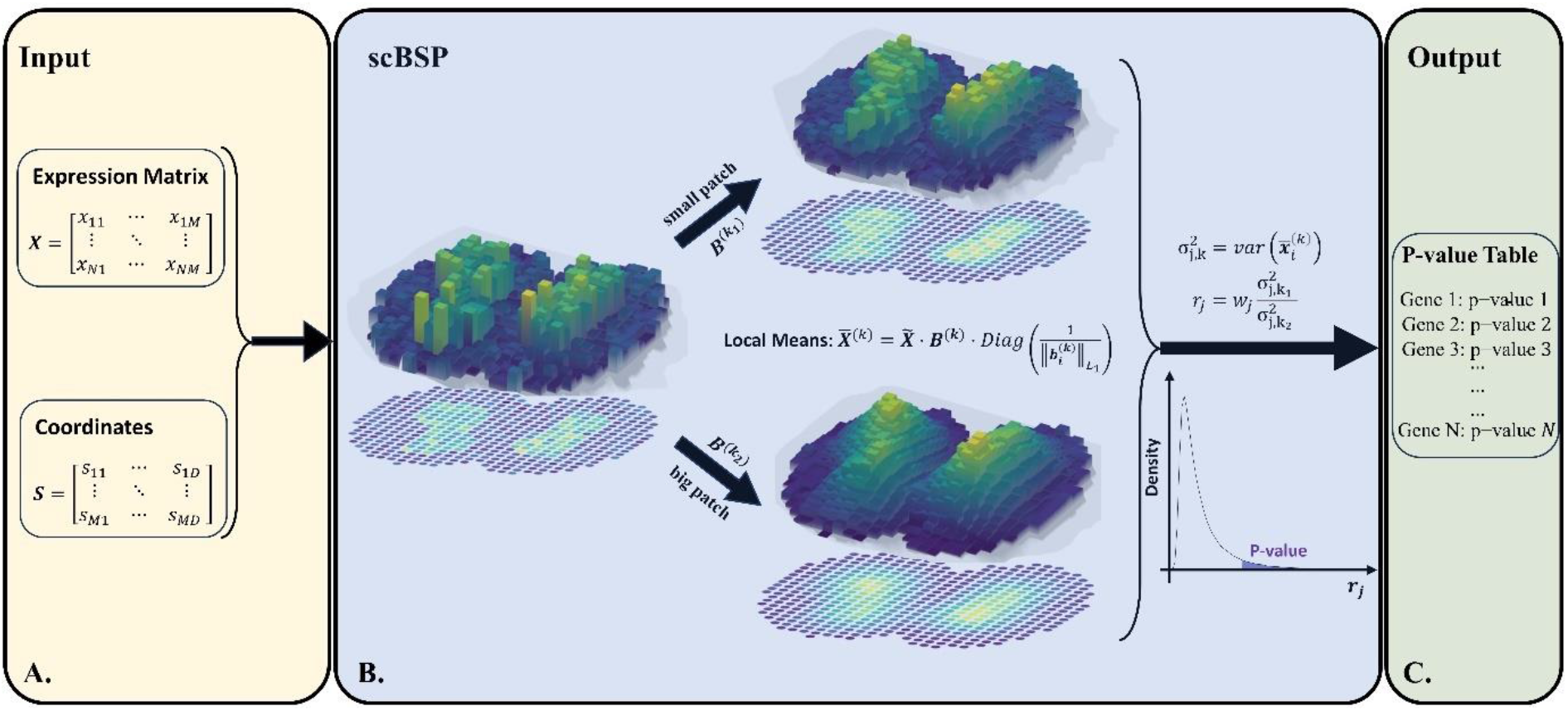
Overview of scBSP. A. Inputs of scBSP. The input comprises an expression matrix, ***X***_***N***×***M***_, containing data for N genes across M spots, and a coordinates matrix, ***S***_*M*×*D*_, representing the spatial coordinates of M spots in a D-dimensional space. B. Schematic of scBSP model. C. Output of scBSP. The output table includes a column of gene names and a column of corresponding p-values.

Given the substantial number of spatial locations and the high sparsity of the expression matrix in large-scale SRT, scBSP adapted the BSP algorithm for expression rescaling, patch determination, and local expression calculation (Supplementary Figure 1) with lower time complexity and space complexity (Table 1). Specifically, for an SRT sample with *M* spots and *N* genes, the *D*-dimensional coordinates of spot *i* are denoted as a *D*-vector ***s***_*i*_ where *i* = 1, …, *M*, and the corresponding coordinates matrix of all spots are denoted as a *M* × *D* matrix 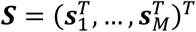. For a spot *i*_0_ in the sample, we define the patch as the set of neighboring spots within the radius *R*_*k*_, denoted as a binary M-vector 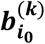 where 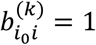 if *dist*(*i*_0_, *i*) < *R*_*k*_ and *i* ≠ *i*. To avoid empty patch, 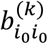 is assigned the value of 1 if and only if 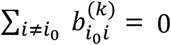. The patches of all spots in the SRT sample are thereby denoted a *M* × *M* binary patch matrix, 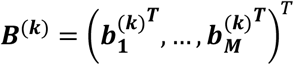, for each given radius *R*_*k*_. We consider a paired big-small patch with a small radius *R*_1_ and *R*_2_, where *R*_1_ < *R*_2_, with the default values as one and three units as demonstrated in BSP (39).

**Table 1:** Time and space complexity of BSP and scBSP. M: number of spots; N: number of genes; S: data sparsity (proportion of non-zero values in the expression matrix).

|  | BSP |  |  | scBSP |  |  |
| --- | --- | --- | --- | --- | --- | --- |
| Process | Method | Time Complexity | Space Complexity | Method | Time Complexity | Space Complexity |
| Data Pre-processing | Min-max Scaling | $O(NM)$ | $O(NM)$ | Maximum Absolute Scaling | $O(M)$ | $O(NM)$ |
| Patch Determination | Exhaustive Search | $O(NM^2)$ | $O(NM)$ | Approximate Nearest Neighbor Search* | $O(NM \log M)$ | $O(NM)$ |
| Expression Matrix Calculation | Dense Matrix | $O(M)$ | $O(NM)$ | Sparse Matrix | $O(SM)$ | $O(SNM)$ |
| Local Expression Calculation | Loop Operation | $O(NM)$ | $O(NM)$ | Matrix Operation | $O(SNM)$ | $O(SNM)$ |

\* The Approximate Nearest Neighbor Search in Python is based on HNSW implementation.

To ensure an adequate number of spots captured by the pre-defined radiuses *R*_1_ and *R*_2_, the coordinates matrix is normalized based on the density of spots such that the average spot-to-spot distance to is slightly less than one unit. The rescaling function is defined as 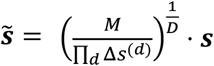, where 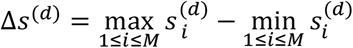 denotes the ranges of coordinates in each direction. The Euclidean distance between spots *i*_1_ and *i*_2_, denoted as *dist*(*i*_1_, *i*_2_), is then calculated as using approximate nearest neighbor search which implements K-Dimensional Tree with the threshold of *R*_*k*_. For the SRT samples with the cell counts ranging from thousands to hundreds of thousands, utilizing the approximate nearest neighbor search can achieve significantly faster running times with a relatively small actual errors compared to the brute-force computation of all distances, as the time complexity is reduced from *O*(*M*^2^) to *O*(*c* × *log*(*M*)), where *c* is a constant depending on the dimension and approximation error (40, 41). The *N* × *M* raw expression matrix is denoted as ***X*** where the raw expression level of gene *j* in spot *i* is denoted as *x*_*ij*_, where 1 ≤ *i* ≤ *M*, 1 ≤ *j* ≤ *N*. Considering the sparsity of high-resolution SRT data, the expressions are rescaled to [0, 1] by genes using the maximum absolute rescaling as

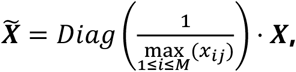

where 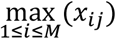 denotes the maximum expression level of gene *j* and 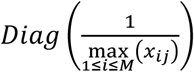 denotes the *N* × *N* diagonal matrix of the *N*-element vector 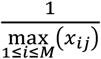. The matrix for averaged expression level 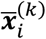 for a given radius *R*_*k*_ at spot *i*, referred to the local means, is calculated as

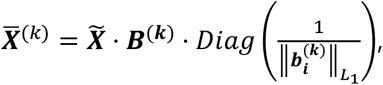

where 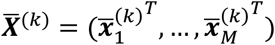 and 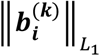 denotes the number of spots in the patch 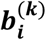. Subsequently, the variance of local means for gene *j* is computed as 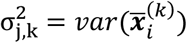. We utilize the ratio between the variances of the paired local averaged expression levels between big patch and small patch, *r*_*j*_, to measure the velocity of changes in the variances of local means for gene *j*, defined as:

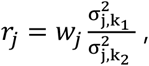

where *w*_*j*_ is the weight to normalize the intrinsic gene expression variance within the gene with the maximum variance, i.e., 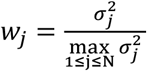, where 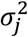 is the variance of raw expression levels of gene *j* of all the spots in the sample, and 1 ≤ *n* ≤ *N*. The distribution of *r*_*j*_ is approximated with a lognormal distribution. The null hypothesis that a gene has no spatial pattern is thus reformulated as the ratio of a gene adhering to the fitted log-normal distribution. To tolerate the potential noise and long-tail deviations, a one-sided p-value is assigned to each gene if *r*_*j*_ exceeds the upper tail of the fitted distribution at a probability of 100 × (1 − *α*)%, where *α* refers to the significance level (usually set as 0.05).

We assume that a minority of genes situated in the upper tail of the distribution exhibit spatial variability, while the majority of genes are non-SVGs. This proposition is particularly relevant in high-throughput SRT platforms like 10X Visium, which deal with an extensive gene repertoire exceeding thousands. Notably, in low-throughput SRT platforms where the gene count is limited (like MERFISH), a set of random genes permuted across spatial locations are recommended to estimate the distribution of the test score for non-SVGs.

### Simulations

#### Simulation for benchmarks

To evaluate the accuracy of identifying SVGs across various scenarios, we conducted model performance assessments of scBSP and compared the statistical power with those obtained using the original BSP algorithm and other existing methods. This evaluation utilized simulated data sourced from previous studies (21, 37–39). The 2D simulations were based on mouse olfactory bulb data, comprising three spatial expression patterns observed across 260 spots (as detailed in the Results section). Each simulation included 1000 simulated SVGs with recognized patterns from studies of SpatialDE and SPARK, alongside 9000 non-SVGs generated via gene permutation without any discernible spatial expression pattern. Statistical power (true positive rates) was calculated using p-values derived from various methods, including basic spatial autocorrelation statistics (e.g., Moran’s I), SpatialDE, SPARK, SPARK-X, SOMDE, BSP, and scBSP, across a wide range of false discovery rates (FDR), accounting for the disparities in the distribution of p-values from each method. Ten replicates were included for each simulation to mitigate sample bias in the power analysis. Two key parameters were considered: signal strengths, quantified as the fold-changes (FCs) between average expressions in patterned and non-patterned regions (FC=3,4,5 for low, moderate, and high signal strengths), and noise level, defined by the dispersion parameters (τ_2_) in SPARK’s model (τ_2_=0.2,0.5,0.8 for low, moderate and high noise levels).

Similar as the 2D simulations, the 3D simulations were composed of 1000 simulated SVGs for each spatial pattern and 9000 permuted genes without any spatial patterns. Three continuous patterns (curved stick, thin plate, and irregular lump) and one discrete pattern (isolated balls) were included (as detailed in the Results section). The simulated data for continuous patterns consisted of 10 segments in the z-coordinate and 225 spots representing cells in each piece in the x- and y-coordinates, assuming that the sample was cryo-sectioned with each section placed on an individual array without any direct contact between array surfaces. The simulations for discrete patterns were designed on the data with 10 segments and 900 spots in each segment, where the x- and y-coordinates on each segment ranging from 0 to 30. The expression of SVGs was sampled based on whether the cell was inside or outside the pattern, distinguishing between marked and non-marked cells. For marked cells inside the pattern, we randomly selected gene expression values from the upper quantile of the gene expression distribution in the seqFISH data. For non-marked cells and those outside the pattern, we assigned gene expression randomly from the expression measurements in the seqFISH data. Non-SVGs were generated by permutating gene expressions of SVGs. Finally, random noise was added proportionally to the averaged standard deviation of expressions in all genes. Three parameters were considered in the simulations including pattern size, measured as the radius of the pattern (radius=1.5,2.0,2.5 for small, moderate, and large pattern size), signal strengths, quantified as the fold-changes between average expressions in patterned and non-patterned regions (FC=2.0,2.5,3.0 for low, moderate, and high signal strengths), and noise level, *σ*, defined as the proportion to the averaged standard deviation of expressions in all genes (*σ*=0,1,2 for low, moderate, and high noise levels). Two additional sets of simulations were incorporated to evaluate the model’s robustness in handling SRT data characterized by distinct inter-plane and within-plane spatial resolutions, as well as the data with varied dropout rates, respectively. To achieve this, the simulated data were generated following the previously described procedure, with an additional step involving the multiplication of z-coordinates by 10 to simulate reduced inter-plane resolution. For the simulations with varied dropout rates, a random subset (10% to 30%) of spots was assigned a value of 0 to simulate the dropout events.

#### Simulations for computational efficiency

Recently studies have shown that SPARK-X is the fastest method for the SVG detection while SOMDE is the second best in most cases but significantly slower compared to SPARK-X (42, 43). However, the results were based on the small samples with the number of spots less than 40, 000. To fairly compare the model performance in the small and large-scale SRT samples respectively, we conducted comparisons of computational efficiency under two distinct scenarios: one involving a larger number of spots with higher data sparsity (referring to HDST data), and the other with a relatively smaller number of spots (referring to stxBrain data).

In the first scenario, two sets of simulations were devised to assess computational efficiency across models. The first set varied the number of spots from 500 to 10,000 while keeping the number of genes fixed at 20,000. The second set varied the number of genes from 2,000 to 40,000 while maintaining a fixed number of spots at 3,000. Expression values were randomly sampled from the expression data of the first anterior of the “stxBrain” dataset in the “SeuratData” package. The metadata of the simulations were detailed in the Supplementary Table 1.

In the second scenario, three sets of simulations were conducted, each varying the number of spots, number of genes, and data density, respectively. The first set involved a range of spots from 50,000 to 400,000, with a fixed number of genes at 20,000 and a data density of 0.05. The second set varied the number of genes from 2,000 to 40,000, with a fixed number of spots at 100,000 and a data density of 0.05. The final set varied the data density from 0.01% to 0.1%, with a fixed number of genes at 20,000 measured across 100,000 spots. This approach was guided by the characteristics of the HDST data, which includes 19,950 genes measured across 181,367 spots, with a data density of 0.04%. Non-zero expression values were generated as follows *Exp*∼⌊1 + 10 × *Beta*(1,5)⌋, where ⌊*x*⌋ denotes the maximum integer that is not greater than x.

### Real Data Analysis

The 10X Genomics Visium mouse brain data were extracted from “stxBrain” data in the “SeuratData” package. For HDST data, we adopted the analysis protocol from SPARK-X. For studies involving Stereo-seq (15), 10x Xenium (44), and Cosmx datasets, we adhered to their respective original protocols. Specifically, for the 10x Xenium and Cosmx datasets, gene expressions were permuted to ensure a total of gene count exceeding 10,000, facilitating sufficient null genes for scBSP and SPARK-X to determine null distributions. The metadata of the simulations were detailed in the Supplementary Table 2. Enrichment analysis was performed using the “clusterProfiler” package in R.

## RESULTS

### scBSP improves computational efficiency by thousandfold on large-scale SRT preserving comparable performance as BSP

We systematically evaluated the computational efficiency of scBSP with seven methods, including Moran’s I, spatialDE, SPARK, SPARK-X, nnSVG, SOMDE, and BSP, for detecting SVGs on simulated SRT data, varying the number of genes and spots (as detailed in the Method section). Considering the high memory usage of some methods (e.g., BSP requires more than 100 GB for some datasets), all methods were executed on a workstation with a 2.40GHz Intel Xeon CPU E5-2680 v4. scBSP outperformed all other tools when analyzing SRT samples with 2000 to 40,000 genes on 3000 spots (Figure 2A). Moreover, scBSP’s running time increased slowly with the number of genes, highlighting the application for SRT data obtained from whole transcriptomic profiling. For SRT samples with 20,000 genes on 500 to 10,000 spots, scBSP also demonstrated faster performance than other tools, particularly with an increase in spot count. While it consumes more memory than SPARK-X, scBSP generally requires less than 2GB of memory usage, making it suitable for processing SRT data with tens of thousands of genes from thousands of spots on personal laptops (Supplementary Figure 2A). Notably, SPARK and nnSVG were terminated manually after running for the maximum allowed time (1 hour) on data with 20,000 genes on more than 500 spots and the data with more than 3,000 genes on 2000 spots (detailed in the Supplementary Table 1), while Moran’s I failed to process any data within the maximum allowed time.

**Figure 2:**
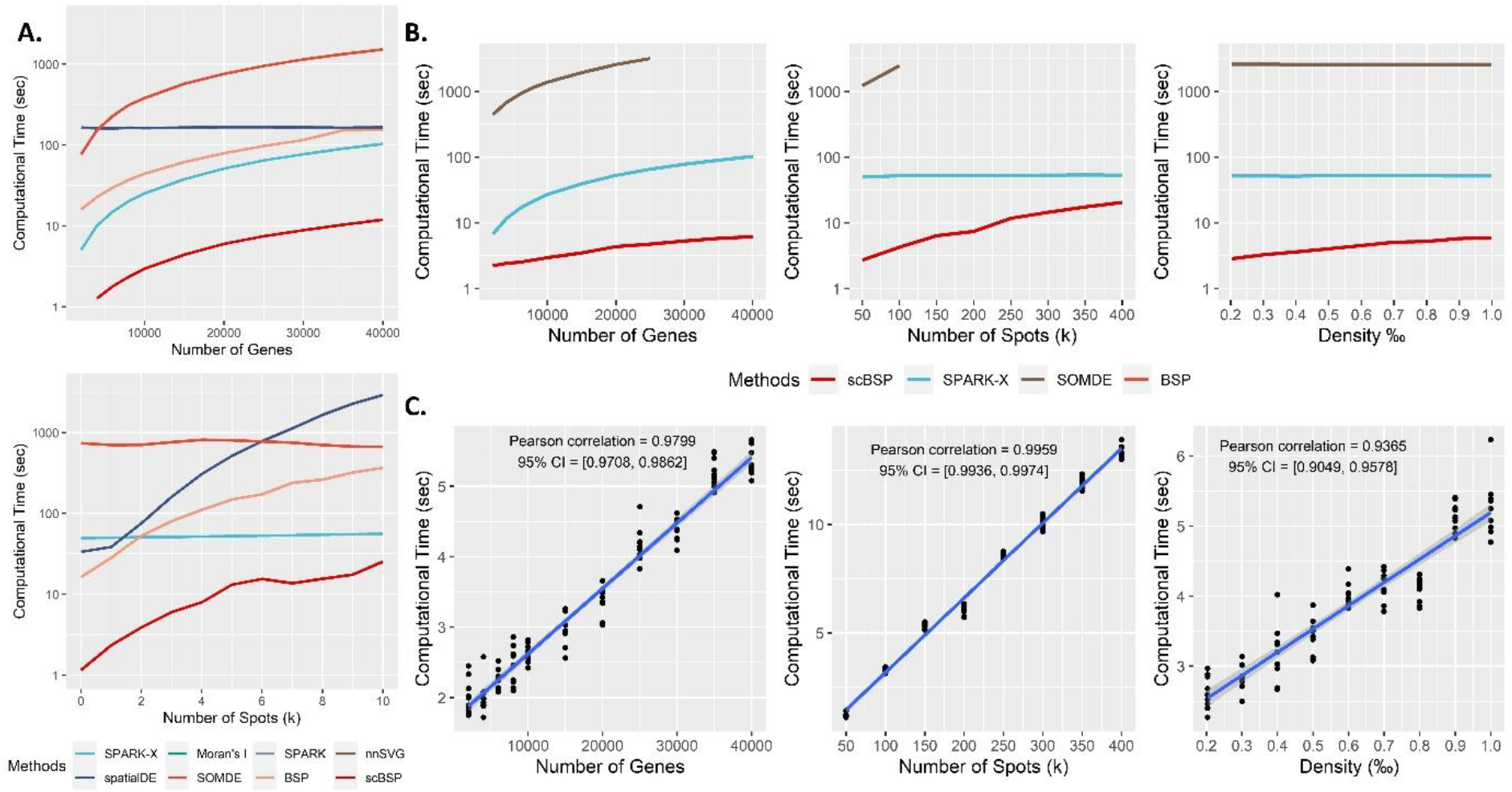
Computational time of scBSP. A. Computational time (y-axis) for analyzing low-resolution SRT data comprising 20,000 genes across 3,000 spots. This analysis varies one parameter while keeping the other constant. B. Computational time (y-axis) for analyzing large-scale SRT data comprising 20,000 genes across 100,000 spots, with a data density of 0.0005. This analysis varies one parameter while keeping the other two constants. C. Computational time (y-axis) of scBSP on the large-scale SRT data (run n = 10 times on a single processor core) with varied gene count, spot count, and data density (x-axis).

We further explored computational efficiency using large-scale simulations comprising 20,000 genes on 100,000 spots with a data density of 0.0005, while varying gene number, spot number, and data density (as detailed in the Method section). scBSP and SPARK-X were the only tools capable of processing such a large sample within 2 hours on a desktop computer. Notably, scBSP analyzed most samples within 20 seconds, outpacing SPARK-X (Figure 2B). Similar to simulations with relatively smaller samples, scBSP’s running time was more sensitive to spot count and data sparsity but less to gene count, highlighting the computational efficiency of scBSP when dealing with large-scale data from whole transcriptome sequencing methods characterized by high expression sparsity. The memory usage of scBSP was higher than SPARK-X but still less than 2GB (Supplementary Figure 2B). scBSP was also tested on mainstream high resolution SRT technologies of 10X Visium, Stereo-seq, HDST, 10X Xenium, and CosMX. Supplementary Table 2 details its running time and memory consumption.

scBSP’s computational efficiency lies in its linear scalability with the number of genes, spots, and data density, which is crucial for detecting SVGs in large-scale SRT data (Figure 2C). To demonstrate this computational linearity, we recorded computational time and peak memory usage when running scBSP on large-scale simulations with 10 replicates. While peak memory usage changes gradually at lower gene counts and data densities due to distance calculation requirements (with 100,000 spots in the large-scale simulations), overall runtime and peak memory usage scale linearly with increasing gene count, spot count, and data density (Supplementary Figure 2C).

To ensure comparable effectiveness to the original BSP algorithm, we applied scBSP to simulations from previous studies, conducting a performance comparison with BSP and other SVG detection methods. Statistical power was assessed using false discovery rate (FDR) to fairly evaluate model performance, considering the discrepancies in the distribution of calibrated p-values across each method. In 2D simulations based on mouse olfactory bulb data with 260 spots (Figure 3A), scBSP exhibited comparable statistical power to BSP across all three spatial patterns (Figure 3B). We also evaluated scBSP’s performance under varied signal strengths (Supplementary Figure 3) and noise levels (Supplementary Figure 4), observing consistent detection capacity in most scenarios, with slight differences in detecting the second spatial pattern with high noise level and the first pattern with low signal strength.

**Figure 3:**
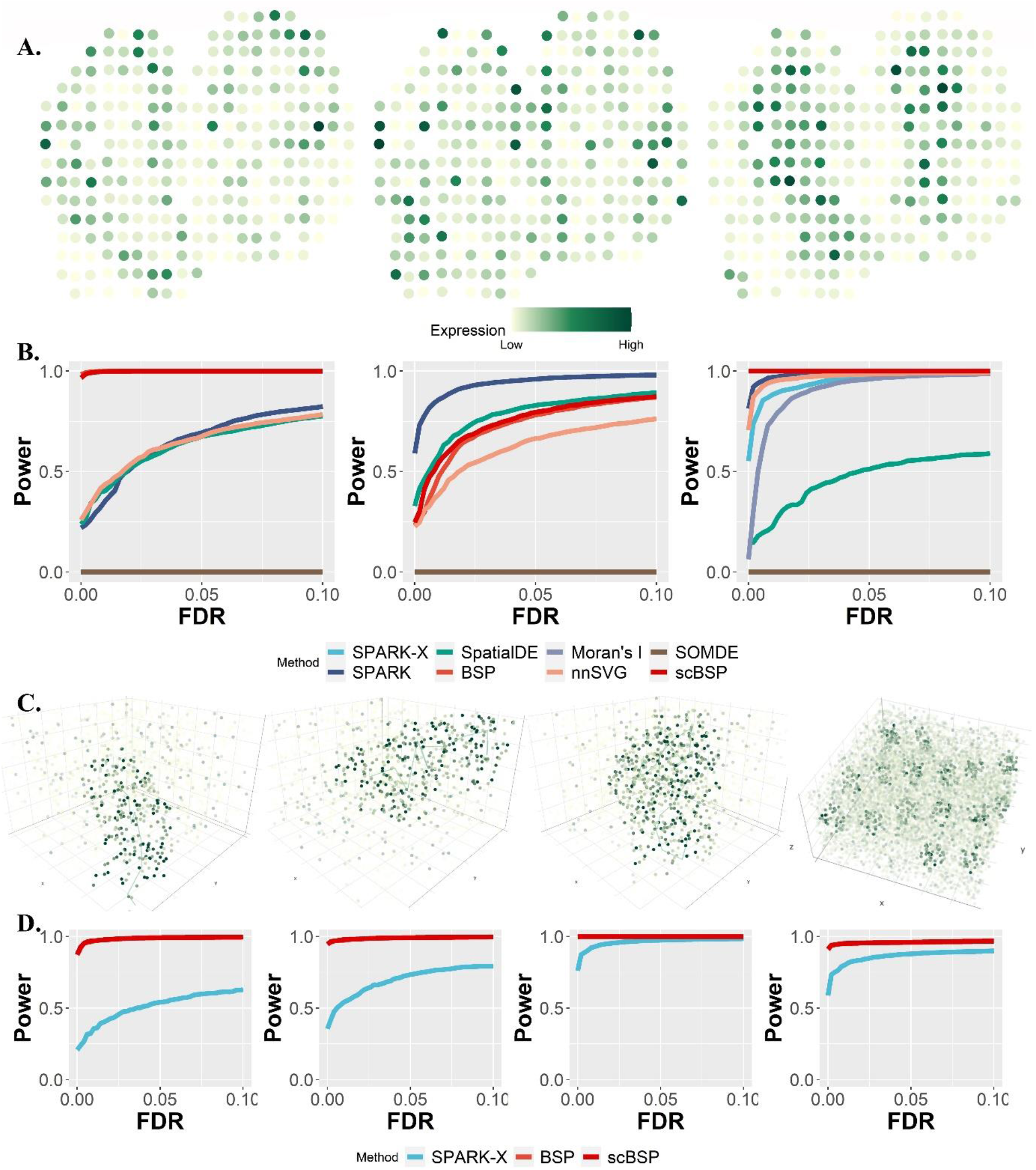
Power analysis on 2D and 3D simulations. A: Spatial patterns I-III in mouse olfactory bulb (left to right) as defined in SpatialDE and SPARK. B: Comparisons of statistical power (y-axis) against false discovery rate (x-axis) on 2D simulations. C: Continuous and discrete spatial patterns in 3D space (left to right). D: Comparisons of statistical power (y-axis) against false discovery rate (x-axis) on 3D simulations.

In 3D simulations with a larger number of spots (ranging from 2000 for continuous spatial patterns to 9000 for discrete patterns), scBSP demonstrated equivalent performance to BSP for all continuous and discrete spatial patterns (Figure 3C), surpassing SPARK-X (Figure 3D). We varied pattern sizes, signal strengths, and noise levels (Supplementary Figure 5-8), and consistently observing robust detection capabilities across scenarios. Additionally, we assessed model performance with varied dropout rates (Supplementary Figure 9) and 3D data featuring inconsistent inter-plane and within-plane spatial resolution (Supplementary Figure 10). From these simulations, scBSP consistently exhibited statistical power comparable to BSP, outperforming SPARK-X in all scenarios.

### scBSP robustly detects SVGs among different SRT technologies

Different SRT technologies reveals the inner characteristics of the tissue samples in different perspectives varying resolution, sensitivity, specificity, and data sparsity. As expression variance on the spatial spaces, SVGs are supposed to be conservative among these different SRT technologies. In addition to the simulations

#### Analysis of high-resolution SRT data from mouse olfactory bulb

To illustrate the practical use of scBSP on large-scale SRT data, we processed three large-scale mouse olfactory bulb data collected from different sequencing techniques (Sections 1 and 2 from Stereo-seq and one from HDST). On a desktop equipped with Intel Core i9-13900 and 32GB memory, scBSP took 53.52 seconds and 65.37 seconds to process the Stereo-seq data with 26,145 genes measured on 107,416 spots and 23,815 genes on 104,931 spots, while SPARK-X took 51.14 seconds and 46.47 seconds, respectively. For the HDST data with more spots and an increased data sparsity, scBSP only took 5.95 seconds to process the whole data which consists of 19,950 genes measured on 181,367 spots, while SPARK-X took 33.16 seconds (Supplementary Table 2).

The detected SVGs from scBSP are consist across the three olfactory bulb datasets from different sequencing technologies from Stero-seq and HDST. Specifically, scBSP identified 2,139 SVGs out of 26,145 genes in the first section from Stereo-seq, 2,057 SVGs out of 23,815 genes in the second section, and 2,292 SVGs out of 19,950 genes in HDST data (p-value<0.05). Overall, scBSP detected a total of 3,080 SVGs, with 1,363 (44.25%) of them shared among all three datasets (Figure 4A). In contrast, only 1.23% were shared over the total number of SVGs identified by SPARK-X. We further assessed the similarity between the identified SVGs from each pair of data using the Jaccard Index (JI). Specifically, for scBSP, the similarity score between two sections of Stereo-seq data was notably high at 0.75, declining to 0.54 and 0.48 when comparing Stereo-seq data with HDST data. In contrast, for SPARK-X, the similarity score was 0.59 between two sections of Stereo-seq data, but decreased rapidly to 0.02 and 0.01 when comparing Stereo-seq data with HDST data.

**Figure 4:**
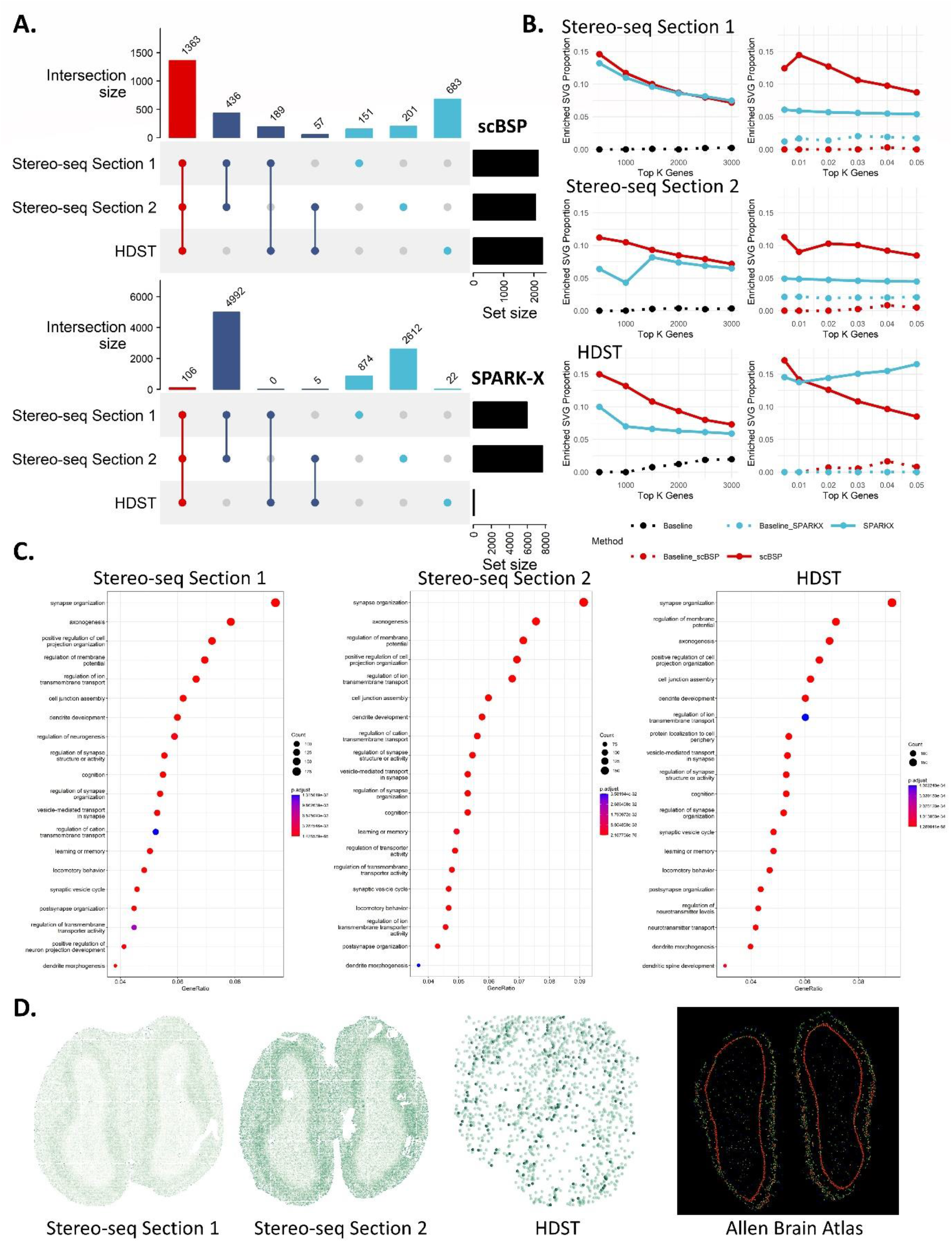
Analysis on mouse olfactory bulb data. A: Upset plot of identified SVGs from scBSP (upper) and SPARK-X (bottom) on three mouse olfactory bulb data. B: Proportions of enriched SVGs over top K genes (left) and proportions of enriched SVGs over the identified SVGs under varied type I error thresholds (right) on each data. C. Enriched gene ontology terms based on scBSP-identified SVGs on each data. D. Expression pattern of gene Ptprd in each dataset and Allen Brain Atlas.

The Gene Ontology (GO) enrichment analysis results exhibited high consistency on scBSP identified SVGs across the three olfactory bulb datasets. The top seven enriched GO terms (synapse organization, axonogenesis, positive regulation of cell projection organization, regulation of membrane potential, regulation of ion transmembrane transport, cell junction assembly, and dendrite development) remained consistent across all three datasets from Stero-seq and HDST (Figure 4C). The olfactory bulb is the primary center in the processing of olfactory information, as it receives, filters, and transmits olfactory signals from the sensory neurons of the olfactory epithelium to one or more cortical olfactory centers, which highly corresponds to the organization of synapses shown in Figure 4C (45, 46). In addition, the olfactory bulb is one of the few neurogenic regions that continues throughout life. This is associated with axonogenesis, dendritic development, and various aspects of learning, and memory performance as illustrated in Figure 4C (47). A representative gene uniquely identified by scBSP, Ptprd (Figure 4D), remained distinguishable even in HDST data characterized by a higher dropout rate (p-values of 6.26e-4, 7.71e-4, and 1.73e-05 for Stereo-seq data section 1, Stereo-seq data section 2, and HDST data, respectively). Ptprd is known to be intricately involved in axon guidance, synapse formation, and cell adhesion within the mouse brain, which has been found to be regulates NMDAR-mediated postsynaptic responses in neural circuits that are spatially linked to asynchronous release sites in recent studies (46, 48, 49). Overall, SVGs identified by scBSP accurately characterized (highly associated with) the biological functions in the mouse olfactory bulb tissue data.

We further investigated the proportions of enriched SVGs relative to the total identified SVGs across varying Type I error thresholds, and their proportions relative to the top K genes (ranging from 500 to 3000) ranked by p-values (Figure 4B). Enriched SVGs were delineated based on enriched GO terms derived from GO enrichment analysis, utilizing either the top K genes with the lowest p-values from scBSP or SPARK-X (left side in Figure 4B), or using the genes with p-values below a specified threshold (right side in Figure 4B). While proportions over the top K genes were similar between scBSP and SPARK-X on Stereo-seq data, proportions based on the Type I error thresholds ranged from 8% to 15% for scBSP, surpassing SPARK-X (4% to 6%). However, proportions over the top K genes from SPARK-X were slightly lower than scBSP in HDST data, which may be resulted from SPARK-X only detected 133 SVGs at a 0.05 threshold. This underscores the accuracy and robustness of scBSP in practical applications, particularly with the utilization of p-value thresholds.

#### Analysis of low-resolution SRT data from whole mouse brain

We further applied scBSP to analyze five whole mouse brain samples with sequencing technologies at a relatively lower resolution. Four sagittal datasets were collected using 10X Genomics Visium platform, while the coronal dataset was obtained from Stereo-seq sequencing. On the desktop, scBSP demonstrated computational efficiency by processing each 10X Visium dataset in an average of 5 seconds, and 25.11 seconds for the Stereo-seq data. In comparison, SPARK-X required 36 seconds per 10X Visium dataset and 47.19 seconds for the Stereo-seq dataset (Figure 5A). Similar to observations in olfactory bulb data, the identified SVGs from scBSP was consistent across various datasets, particularly exhibiting high consistency between the two anterior sections (JI=0.92) and two posterior sections (JI=0.94) (Figure 5B). Moreover, the enriched pathways detected by scBSP were very close across datasets, which were related to the biological processes in mouse brain (Supplementary Figure 11-15). We also examined the proportions of enriched SVGs over the identified SVGs across varied type I error thresholds (Figure 5C). The threshold-based proportion of scBSP was consistently higher than SPARK-X, which highlights the application of scBSP as the SVGs were typically selected based on a given threshold of type I error rate in practice. In summary, scBSP efficiently and robustly identified biological meaningful SVGs in SRT invariant to the sequencing technologies and resolutions.

**Figure 5:**
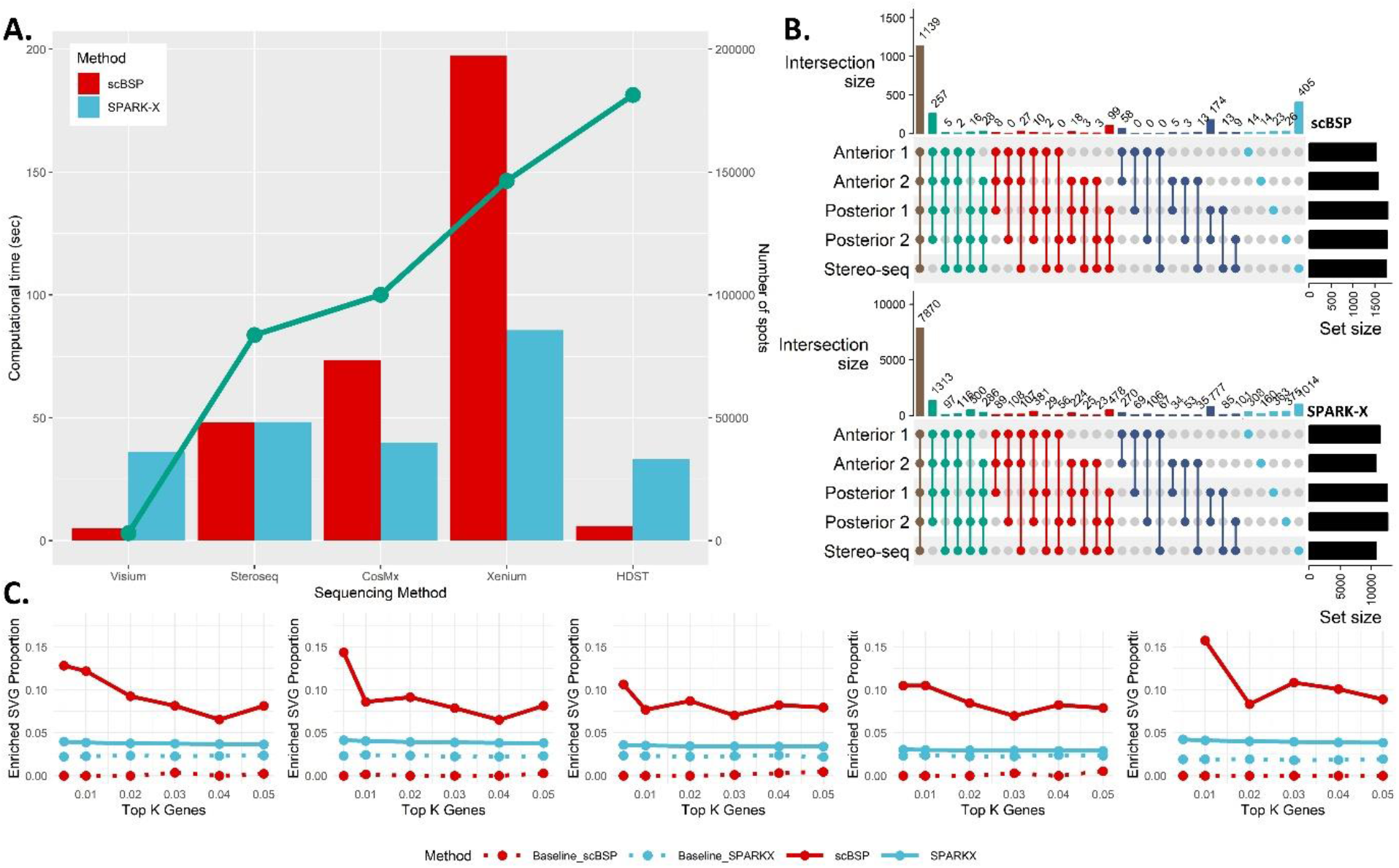
Analysis on low-resolution mouse brain data. A. Computational time of scBSP and SPARK-X on real data. The bar charts illustrated running time (left y-axis) on each data (x-axis). The green line denotes number of spots in each dataset (right y-axis). B: Upset plot of identified SVGs from scBSP (upper) and SPARK-X (bottom) on five mouse brain data. C: Proportions of enriched SVGs over top K genes (left) and proportions of enriched SVGs over the identified SVGs under varied type I error thresholds (right) on each data.

## DISCUSSION

In this study, we introduced scBSP, a novel computational efficiency package available in both R and Python, designed for identifying SVGs in large-scale SRT data. scBSP implements sparse matrix operation to takes advantages of algorithm optimization and compiler optimization in contemporary computer architectures. The scBSP R package is conveniently hosted on the Comprehensive R Archive Network (CRAN), requiring no external software dependencies other than R packages, ensuring straightforward installation and usability across all major operating systems. Similarly, the scBSP Python library is readily downloadable and installable via the Python Package Index (PyPI), offering both a full version reliant on Microsoft Visual C++ for approximate nearest neighbor search and a lightweight version with no dependencies beyond Python libraries. Additionally, the scBSP R package boasts a high level of interoperability with other tools, accepting inputs in the form of Seurat objects, thus seamlessly integrating with a wide array of analysis and visualization functions within the Bioconductor ecosystem.

scBSP addresses the pressing need for tools capable of detecting SVGs in large-scale SRT data, particularly those characterized by high sparsity. The advent of high-resolution spatial sequencing techniques has seen a drastic increase in the number of spots per SRT sample, escalating from hundreds to millions over the past decade. This surge in resolution poses a challenge for most of the existing tools, many of which require hours to days to process a single SRT sample. In contrast, scBSP demonstrates remarkable efficiency, processing samples with hundreds of thousands of spots in mere seconds to minutes. For instance, on a typical desktop, scBSP processed Stereo-seq data containing 107,416 spots in just 53.52 seconds, presenting a comparable alternative to SPARK-X in R. Notably, the scBSP Python library stands as the only tool available for handling such large-scale data within reasonable computational resources. This improved computational efficiency not only accelerates SVG inference in Python but also makes spatial transcriptomics accessible to more researchers, driving faster insights into spatial gene expression patterns across various biological processes and allowing for the implementation of well-developed deep learning frameworks among the Python community. Moreover, when analyzing data from HDST sequencing comprising 181,367 spots, scBSP took 5.95 seconds on the desktop. Compared to the Stereo-seq data, the computational time decreases despite an increase in spot count, since the data density decreases from 2% to 0.04%. This further highlights the potentials of scBSP in the high-resolution SRT data where the expression data is usually highly sparse.

Furthermore, scBSP’s non-parametric, spatial granularity-based model offers a fast, accurate, and robust solution for SVG detection in large-scale SRT data. Simulations demonstrate scBSP’s superior performance in most scenarios, exhibiting the robustness to varying signal strengths, noise levels, and dropout rates. Moreover, when applied to large-scale olfactory bulb datasets collected from different sequencing techniques, the identified SVGs from Stereo-seq and HDST data exhibit high consistency (JI = 0.54) despite a significant increase in the data sparsity, while the number of identified SVGs from SPARK-X decreased from 5972 to 133 (JI = 0.02). This is potentially because the spatial granularity-based model does not make assumptions about the underlying distributions of expressions, which may not hold in the complex biological environment and the expression data with high dropout rates. This comparison across SRT samples highlights scBSP’s reliability and robustness, essential for obtaining meaningful insights across experiments and sequencing technologies. Notably, this characteristic of scBSP will benefit more with the advancement of new high-throughput, high-sensitivity sequencing technologies in the future. Additionally, scBSP’s ability to detect enriched SVGs and their consistent enrichment in gene ontology terms across datasets further validates its effectiveness in capturing biologically relevant information.

While the scBSP package offers significant advancements in SVG detection on large-scale SRT data, it still presents some limitations. Unlike its R package, the scBSP library in Python relies on external software (Microsoft Visual C++ 14 or higher) to detect neighboring spots within a specified radius using the nearest neighbor search method, as equivalent functions are lacking. To address this issue, we offer a lightweight version utilizing Ball Tree as an alternative method for patch determination. However, this may result in lower computational efficiency, particularly with larger spot counts, and could yield minor discrepancies in results compared to the R implementation, owing to methodological differences in distance calculation. Additionally, scBSP’s computational time and memory usage are sensitive to data sparsity in large-scale SRT data. While it outperforms SPARK-X on the benchmarks, it may require more time and memory than SPARK-X for large-scale data with low sparsity, such as Xenium and CosMx SRT data where genes are pre-selected.

In the future, we will delve into scBSP’s applications across a range of high-resolution SRT case studies, including analyses of the tumor microenvironment, Alzheimer’s disease, and kidney research. We also aim to explore scBSP’s potential with other spatial omics data at the cellular or subcellular level, such as spatial CITE-seq and spatial CUT&Run-seq. Furthermore, we plan to investigate SVGs in large-scale spatial tempo studies, like embryogenesis in mouse embryos using Stereo-seq platforms.

In conclusion, scBSP presents as a versatile and user-friendly package available in both R and Python, facilitating SVG detection on large-scale SRT data. It offers accurate and robust SVG detection across SRT data with varying spatial resolutions, while the computational time scales linearly with cell counts, gene counts, and data sparsity. This makes it particularly well-suited for analyzing large-scale SRT data, especially data from whole transcriptomic profiling with high sparsity. As high-resolution spatial sequencing techniques continue to evolve rapidly, scBSP’s scalability, robustness, and precision in capturing biologically relevant information render it an invaluable tool for researchers investigating spatial gene expression patterns and their implications across diverse biological processes and diseases.

## DATA AVAILABILITY

All relevant data supporting the key findings of this study are available within the article and its Supplementary Information files. The stxbrain data can be downloaded with “SeuratData” package in R. HDST data are available at Broad Institute’s single-cell repository with ID SCP420. Stereo-seq data from the Mouse Organogenesis Spatiotemporal Transcriptomic Atlas (MOSTA) at https://db.cngb.org/stomics/mosta/download/. 10x Xenium human breast cancer data and mouse brain data are available at https://www.10xgenomics.com/products/xenium-in-situ/preview-dataset, and https://www.10xgenomics.com/datasets/fresh-frozen-mouse-brain-replicates-1-standard.

Cosmx dataset (Lung Cancer Slices) are available at https://nanostring.com/products/cosmx-spatial-molecular-imager/ffpe-dataset/nsclc-ffpe-dataset/. The simulation data are available in Figshare database at https://doi.org/10.6084/m9.figshare.2418792360.

## Code availability

The Python library, “scBSP”, is available at https://pypi.org/project/scbsp/ and a corresponding R package, “scBSP”, is also available on R CRAN at https://cran.r-project.org/web/packages/scBSP/index.html. Project home page: https://github.com/CastleLi/scBSP/ https://github.com/YQ-Wang/scBSP Archived versions: https://zenodo.org/records/11123268

## AUTHOR CONTRIBUTIONS

Conceptualization: JL, JW, QM, and DX; methodology: JL, and JW; software coding: JL and YW; data collection and investigation: JL, JW, and MR; data analysis: JL, YW, LS, CX and MR; pathology analysis: QG, and YC; software testing and tutorial: JL, and YW; manuscript writing, review, and editing: JL, JW, QM, and DX.

