## Supplementary Figures for "scBSP: A fast and accurate tool for identifying spatially variable genes from spatial transcriptomic data"

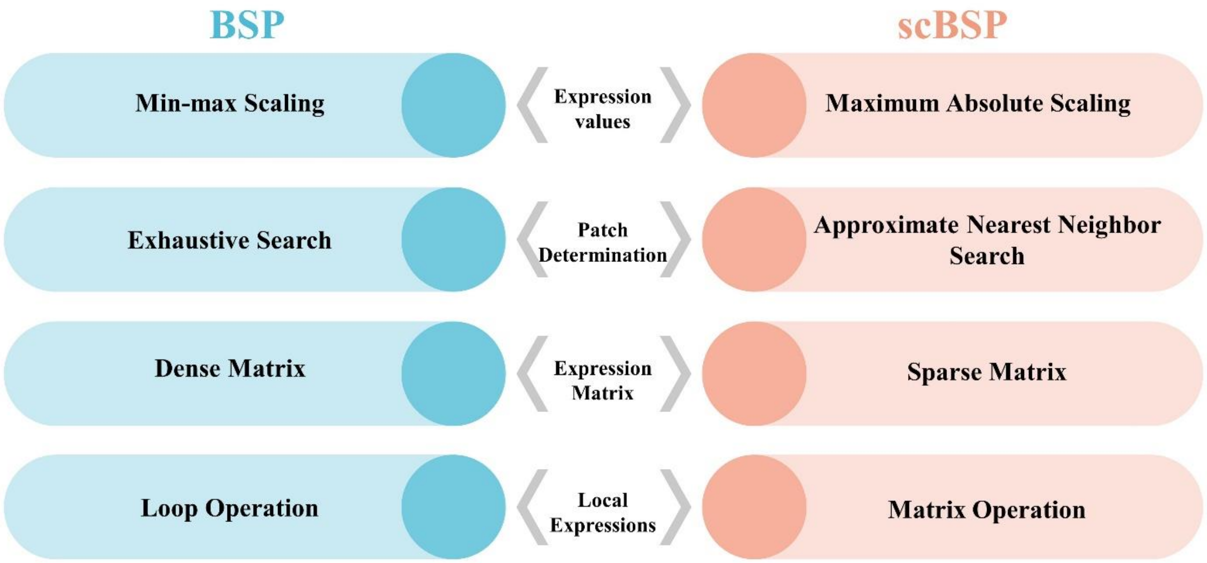


Supplementary Figure 1. Comparisons of scBSP and BSP.

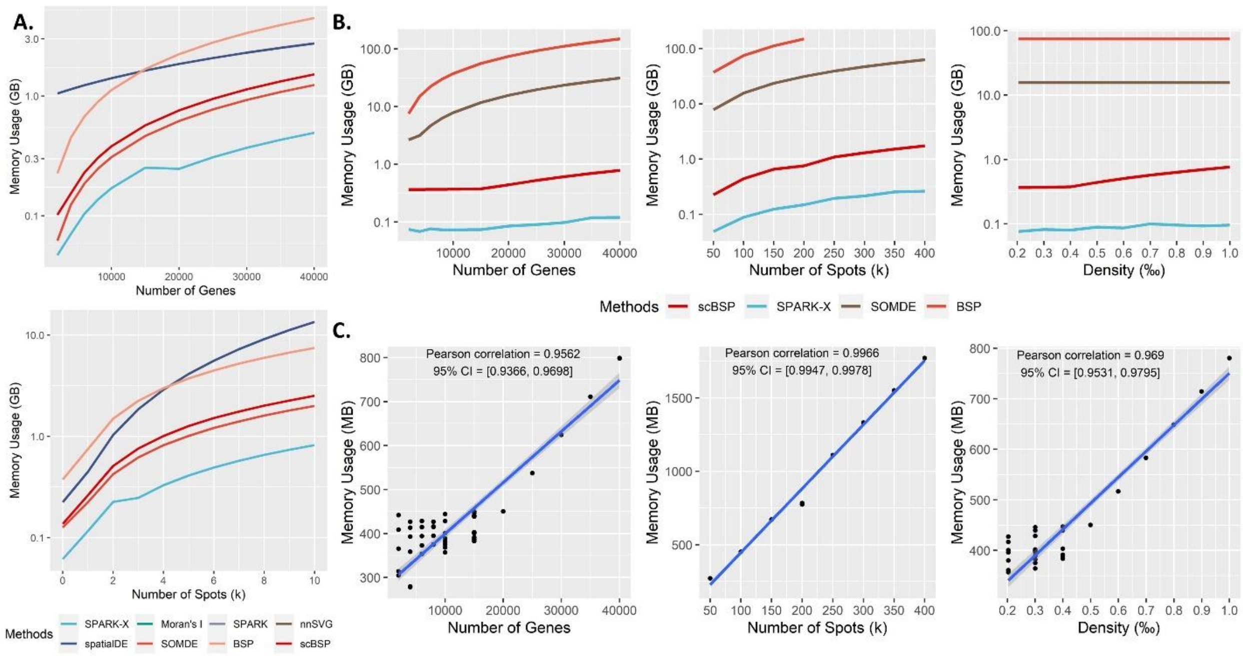


Supplementary Figure 2. A: Memory usage (y-axis) for analyzing SRT data comprising 20,000 genes across 3,000 spots. B: Memory usage (y-axis) for analyzing large-scale SRT data comprising 20,000 genes across 100,000 spots, with a data density of 0.0005. This analysis varies one parameter while keeping the other two constants. C: Memory usage (y-axis) of scBSP on the large-scale SRT data (run n = 10 times on a single processor core) with varied gene count, spot count, and data density (x-axis).

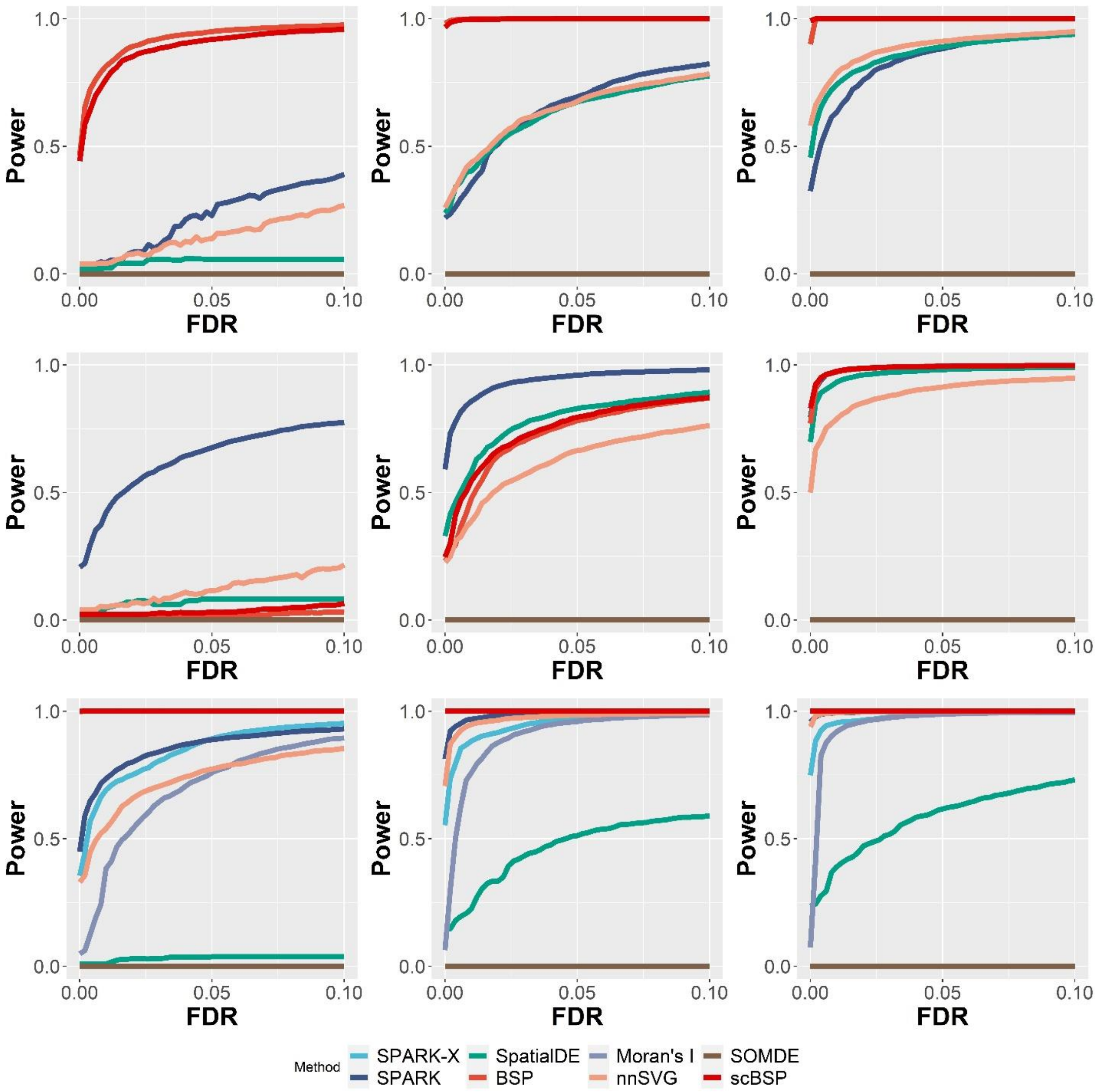


Supplementary Figure 3. Statistical power on 2D simulations with varied signal strengths. Signal strengths were measured as the fold-changes in the averaged expressions between the pattern and non-pattern regions. Power curves were drawn using the averaged statistical power (y-axis) across ten replicates against the false discovery rates (x-axis) for the detected SVGs from each method. Results with weak (FC = 3), moderate (FC = 4), and high (FC = 5) signal strengths were shown in the left, middle, and right columns, while the upper, middle, and bottom rows represented the results from three spatial patterns in Fig.2 A. All simulations were generated using a fixed moderate noise level.

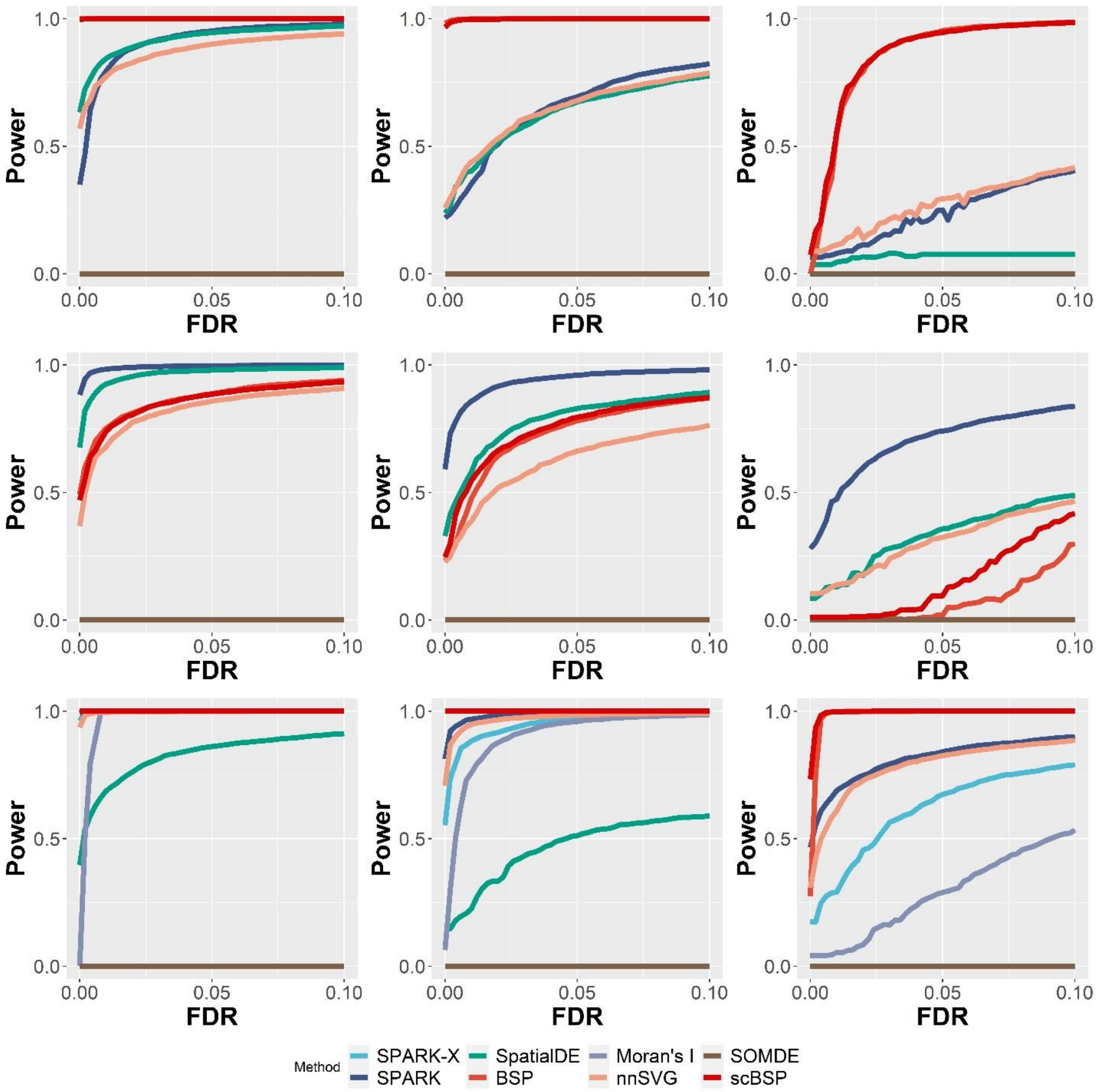


Supplementary Figure 4. Statistical power on 2D simulations with varied noise levels. Noise levels were defined as the dispersion parameters ($\tau_2$) in SPARK's model. Power curves were drawn using the averaged statistical power (y-axis) across ten replicates against the false discovery rates (x-axis) for the detected SVGs from each method. Results with low ($\tau_2$=0.2), moderate ($\tau_2$=0.5), and high ($\tau_2$=0.8) noise levels were shown in the left, middle, and right columns, while the upper, middle, and bottom rows represented the results from three spatial patterns in Fig.2 A. All simulations were generated using a fixed moderate signal strength.

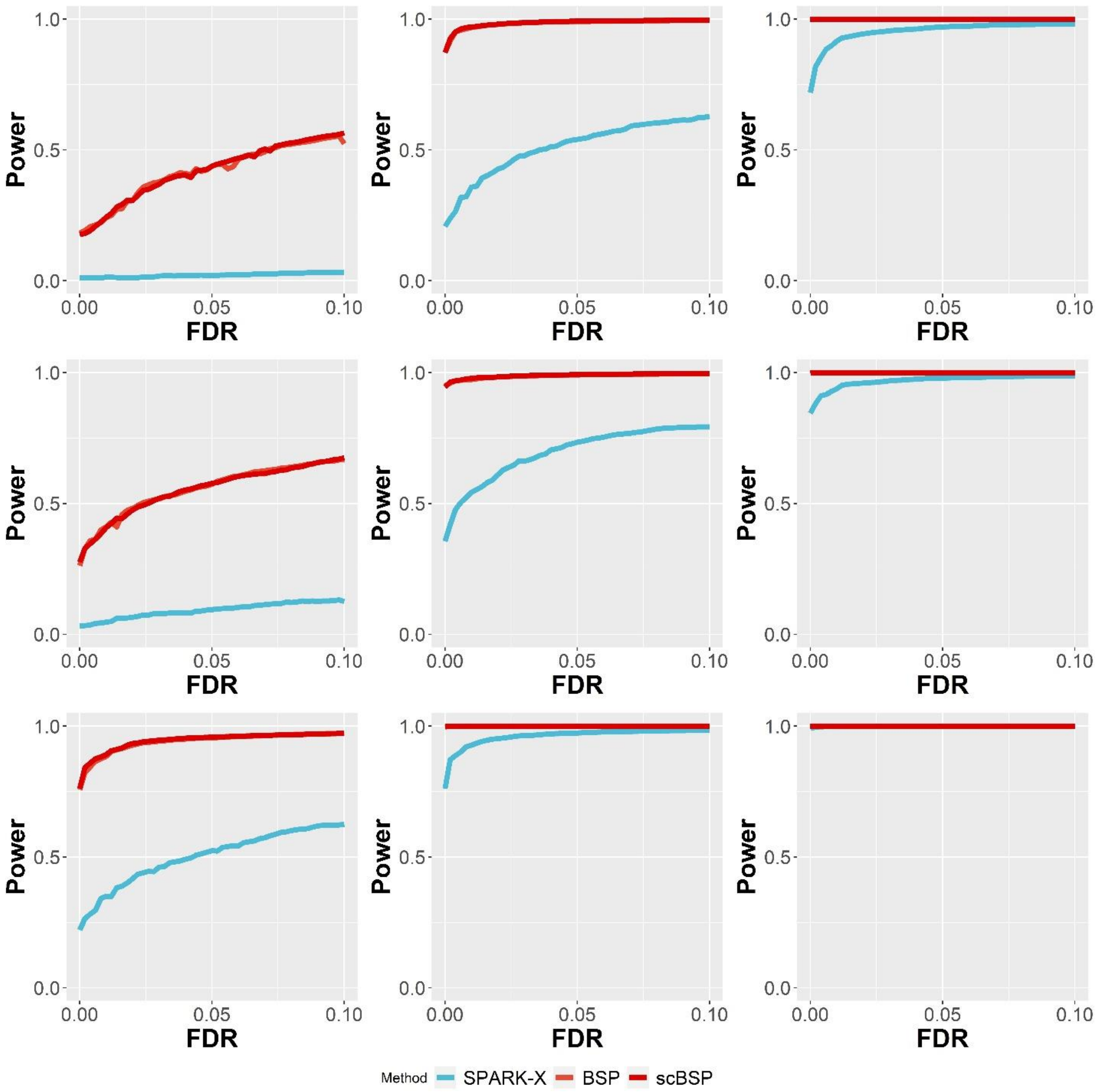


Supplementary Figure 5. Statistical power on 3D simulations of continuous patterns with varied pattern size. Pattern size was measured as the radius of the pattern as described in the Method section. Power curves were drawn using the averaged statistical power (y-axis) across ten replicates against the false discovery rates (x-axis) for the detected SVGs from each method. Results with small (radius=1.5), moderate (radius=2.0), and large (radius=2.5) pattern sizes were shown in the left, middle, and right columns, while the upper, middle, and bottom rows represented the results from three continuous spatial patterns in Fig.2 C. All simulations were generated using a fixed moderate signal strength and noise level.

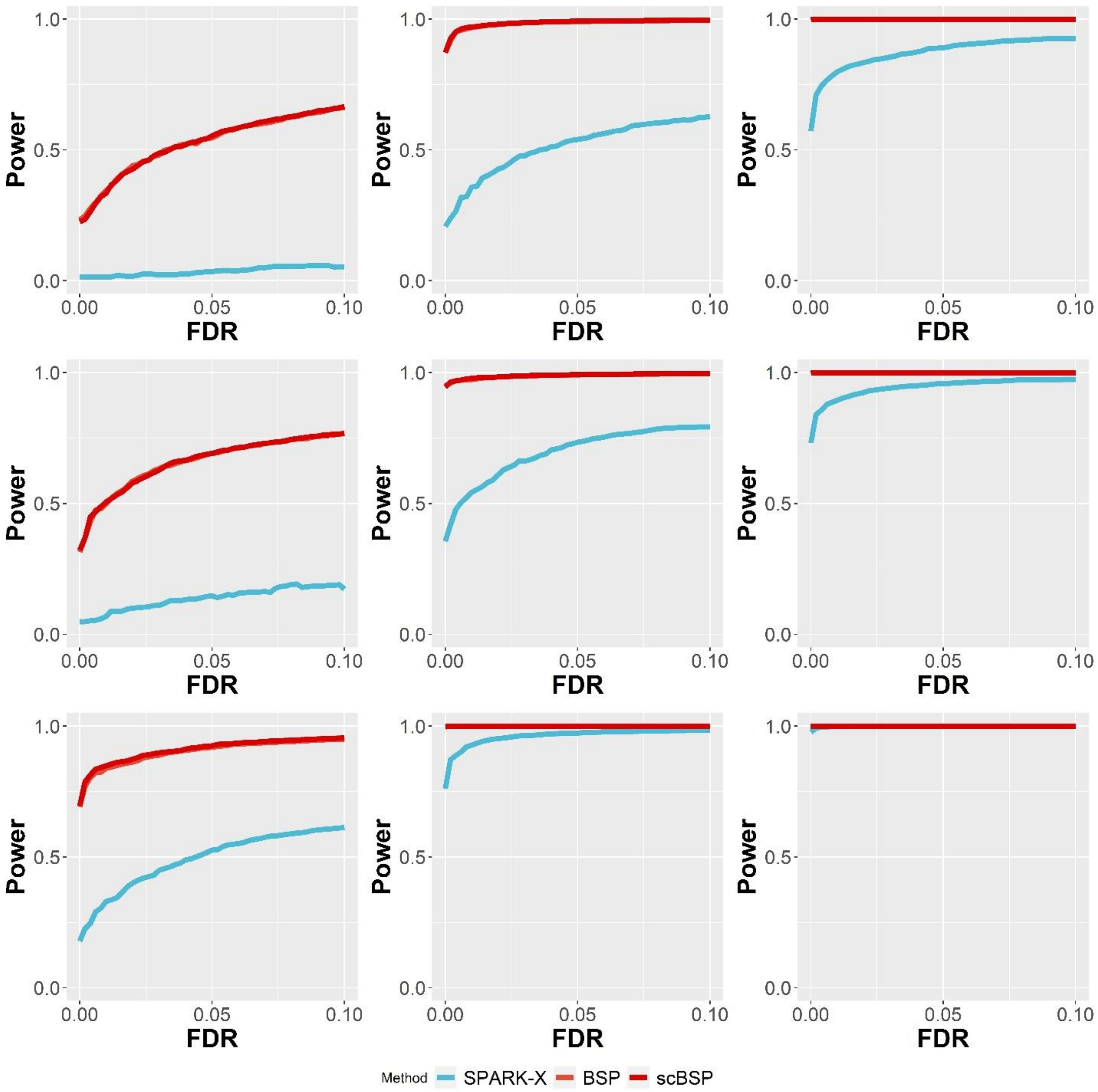


Supplementary Figure 6. Statistical power on 3D simulations of continuous patterns with varied signal strengths. Signal strengths were measured as the fold-changes in the averaged expressions between the pattern and non-pattern regions. Power curves were drawn using the averaged statistical power (y-axis) across ten replicates against the false discovery rates (x-axis) for the detected SVGs from each method. Results with weak (FC = 2.0), moderate (FC = 2.5), and high (FC = 3.0) signal strengths were shown in the left, middle, and right columns, while the upper, middle, and bottom rows represented the results from three continuous spatial patterns in Fig.2 C. All simulations were generated using a fixed moderate pattern size and noise level.

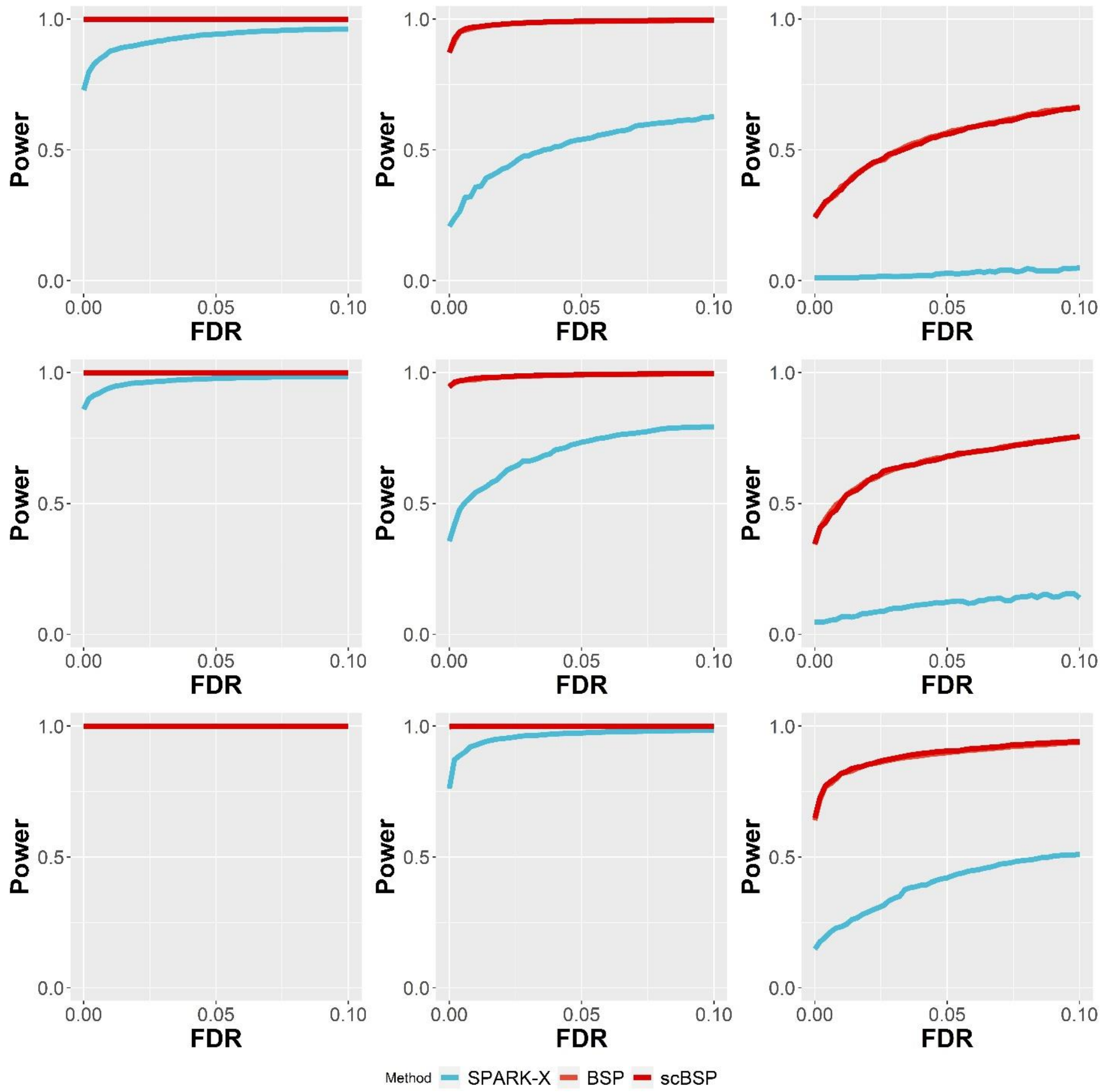


Supplementary Figure 7. Statistical power on 3D simulations of continuous patterns with varied noise levels. Noise levels ($\tau$) were measured as the proportions to the averaged standard deviation of simulated genes (detailed in the Method section). Power curves were drawn using the averaged statistical power (y-axis) across ten replicates against the false discovery rates (x-axis) for the detected SVGs from each method. Results with low ($\tau = 0$), moderate ($\tau = 1$), and high ($\tau = 2$) noise levels were shown in the left, middle, and right columns, while the upper, middle, and bottom rows represented the results from three continuous spatial patterns in Fig.2 C. All simulations were generated using a fixed moderate pattern size and signal strength.

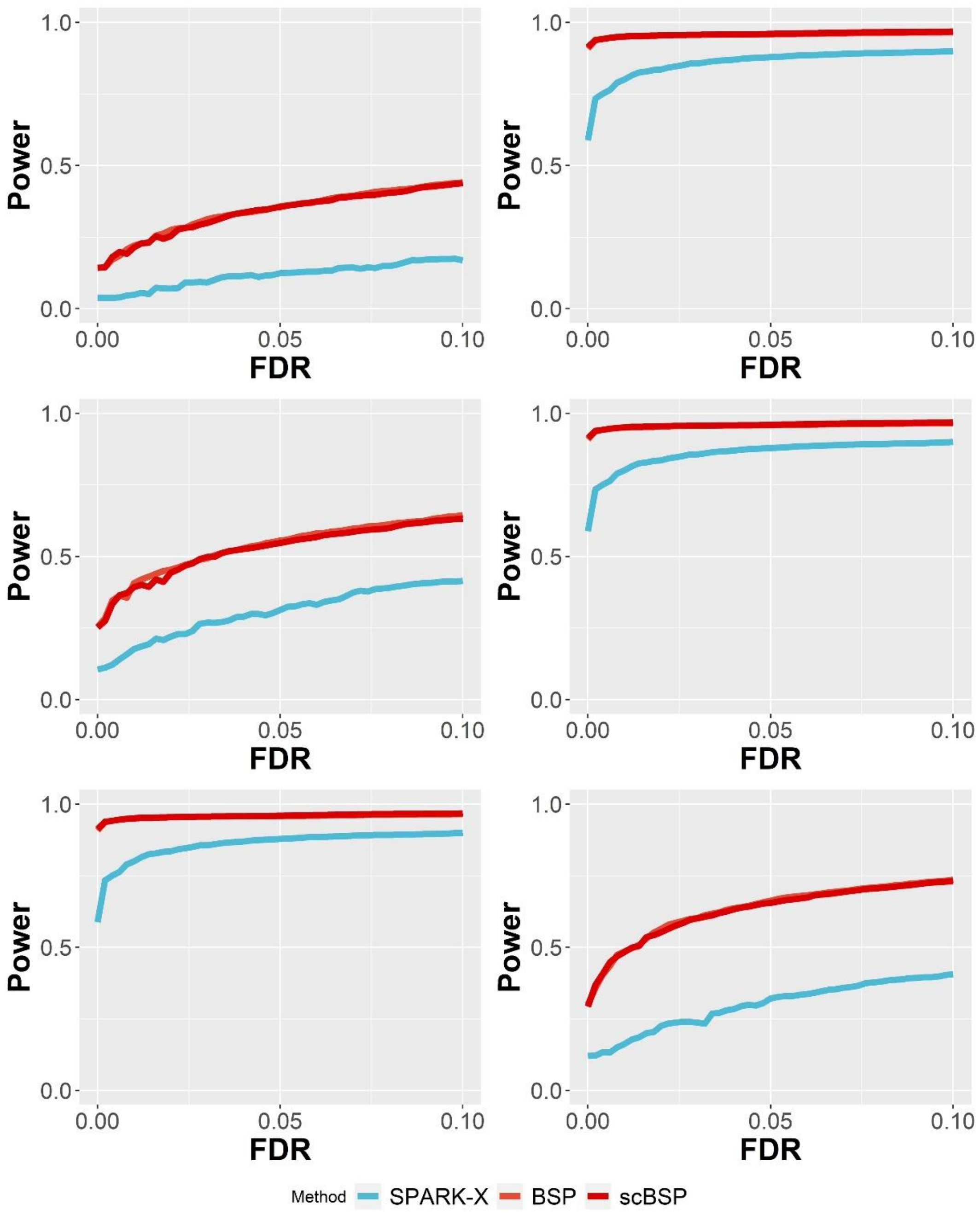


Supplementary Figure 8. Statistical power on 3D simulations of discrete patterns. Power curves were drawn using the averaged statistical power (y-axis) across ten replicates against the false discovery rates (x-axis) for the detected SVGs from each method. Results with varied pattern size (left: radius=1.5; right: radius=2.0), signal strengths (left: FC=2.0; right: FC=2.5) and noise levels (left: $\tau = 2.0$; right: $\tau = 3.0$) were shown in the upper, middle, and bottom rows as described in the Method section.

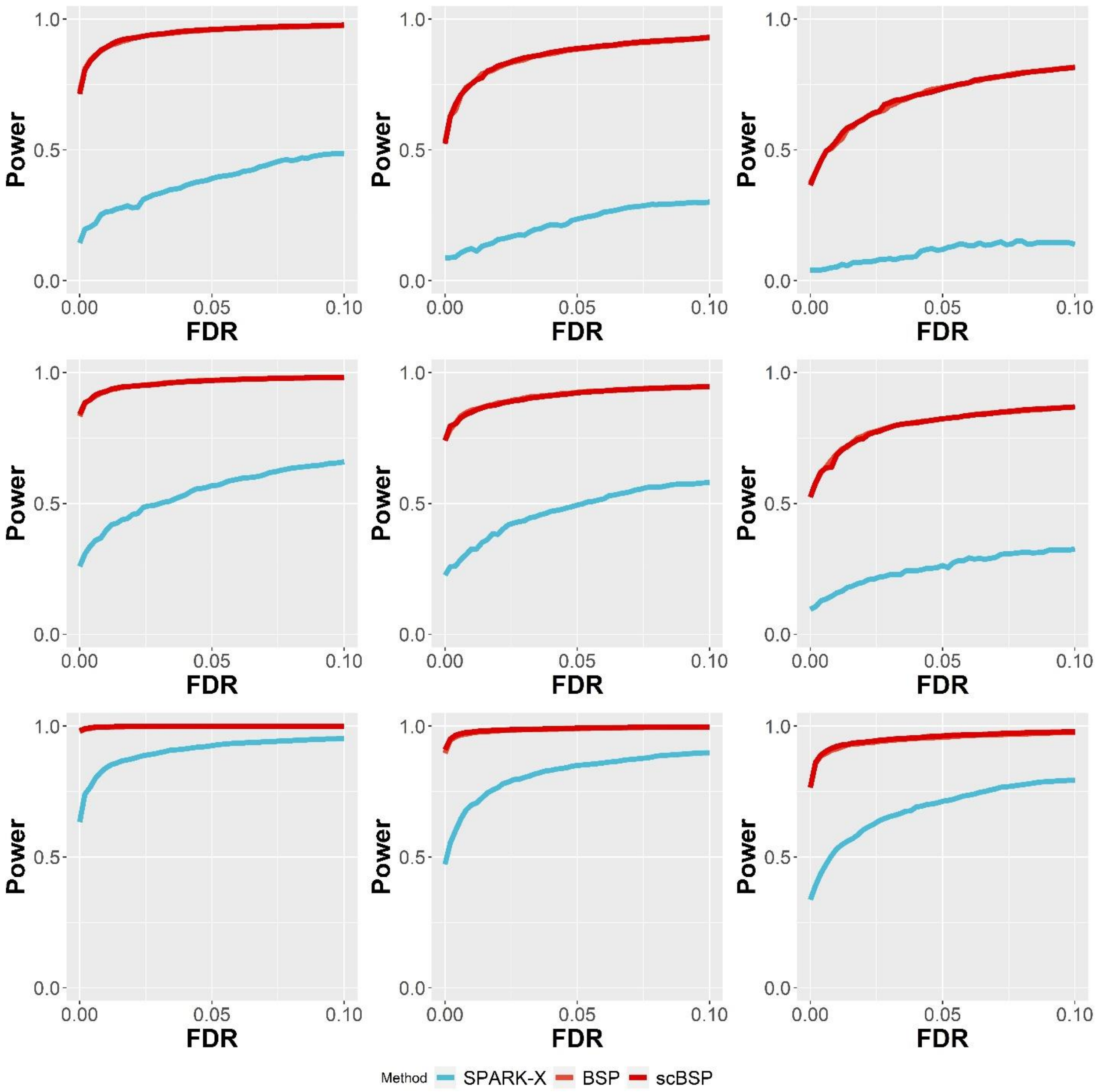


Supplementary Figure 9. Statistical power on 3D simulations of continuous patterns with varied dropout rates. Power curves were drawn using the averaged statistical power (y-axis) across ten replicates against the false discovery rates (x-axis) for the detected SVGs from each method. Results with low (10%), moderate (20%), and high (30%) dropout rates were shown in the left, middle, and right columns, while the upper, middle, and bottom rows represented the results from three continuous spatial patterns in Fig.2 C. All simulations were generated using a fixed moderate pattern size and signal strength.

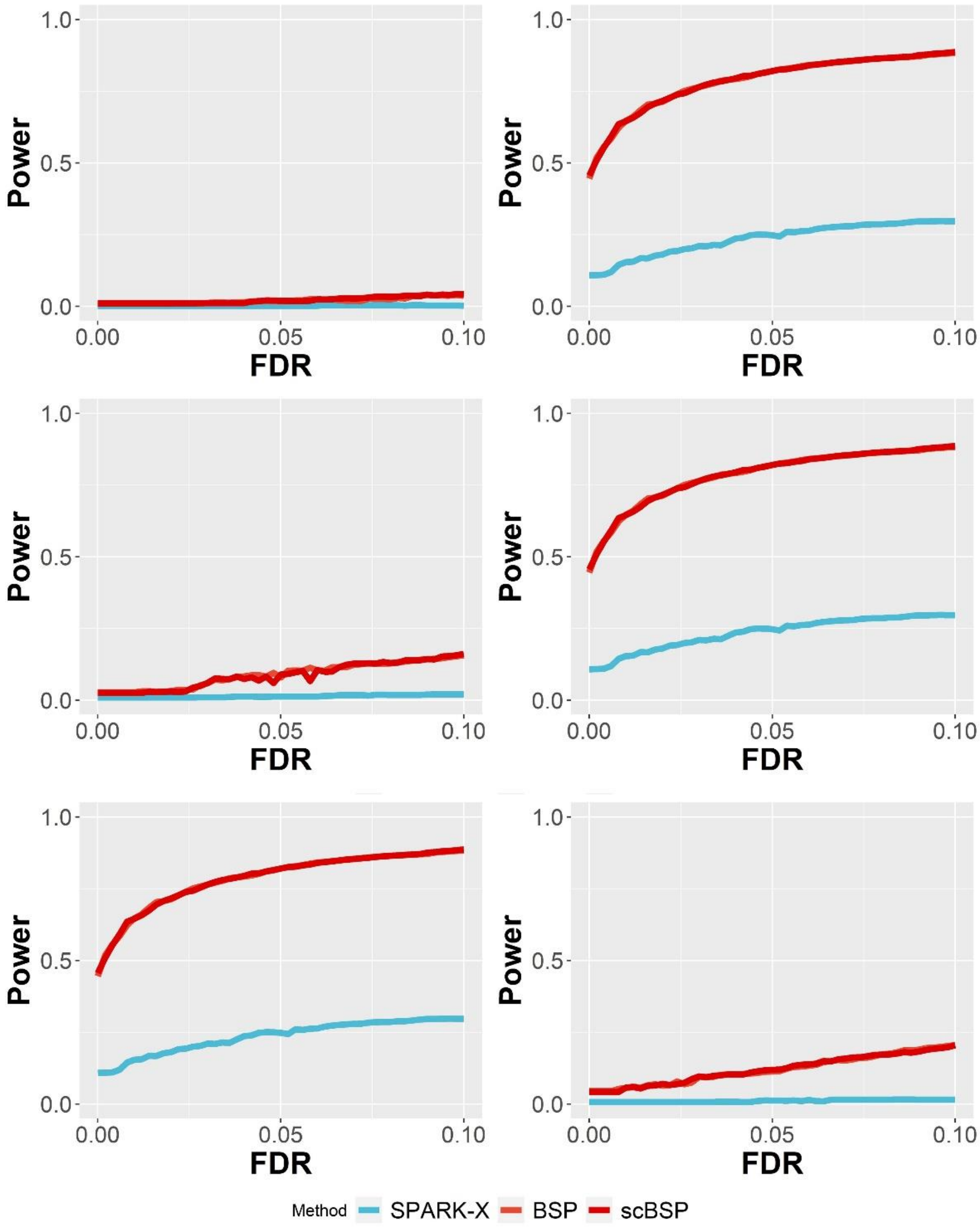


Supplementary Figure 10. Statistical power on 3D simulations of with inconsistent inter-plane and within-plane spatial resolution. Power curves were drawn using the averaged statistical power (y-axis) across ten replicates against the false discovery rates (x-axis) for the detected SVGs from each method. Results with varied pattern size (left: radius=1.5; right: radius=2.0), signal strengths (left: FC=2.0; right: FC=2.5) and noise levels (left: $\tau = 1.0$; right: $\tau = 2.0$) were shown in the upper, middle, and bottom rows as described in the Method section.

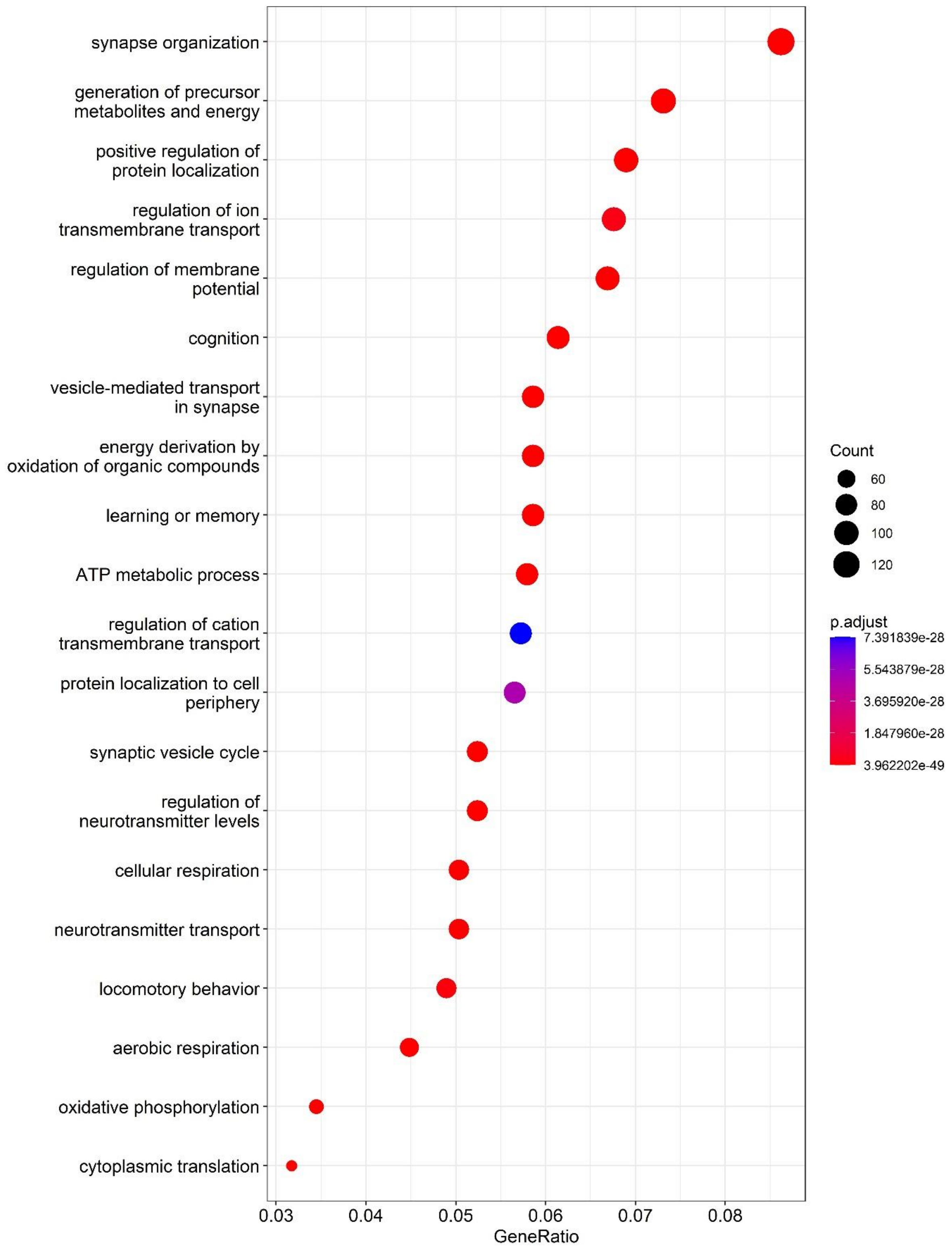


Supplementary Figure 11. Enriched gene ontology terms on 10X Visium mouse brain anterior 1 data.

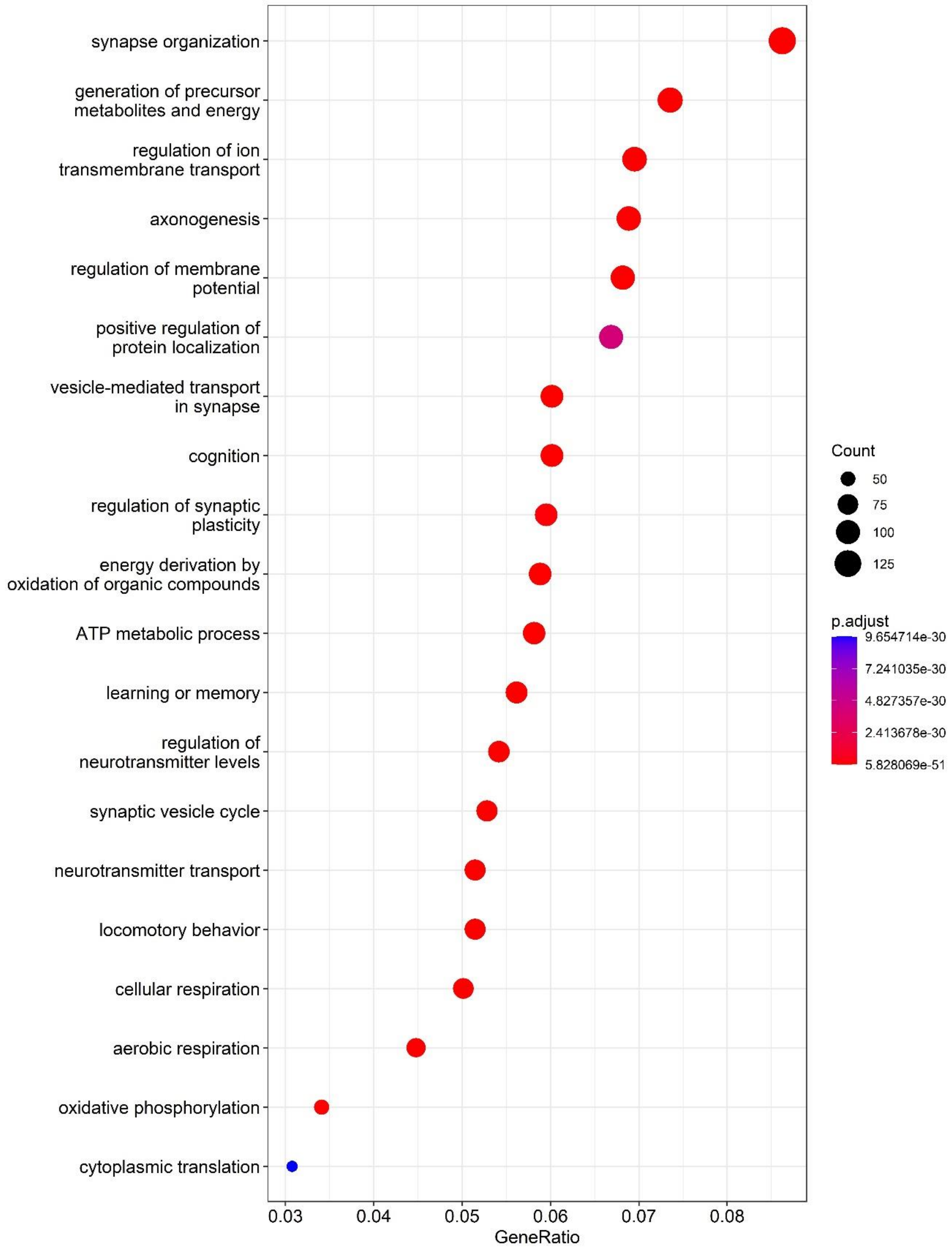


Supplementary Figure 12. Enriched gene ontology terms on 10X Visium mouse brain anterior 2 data.

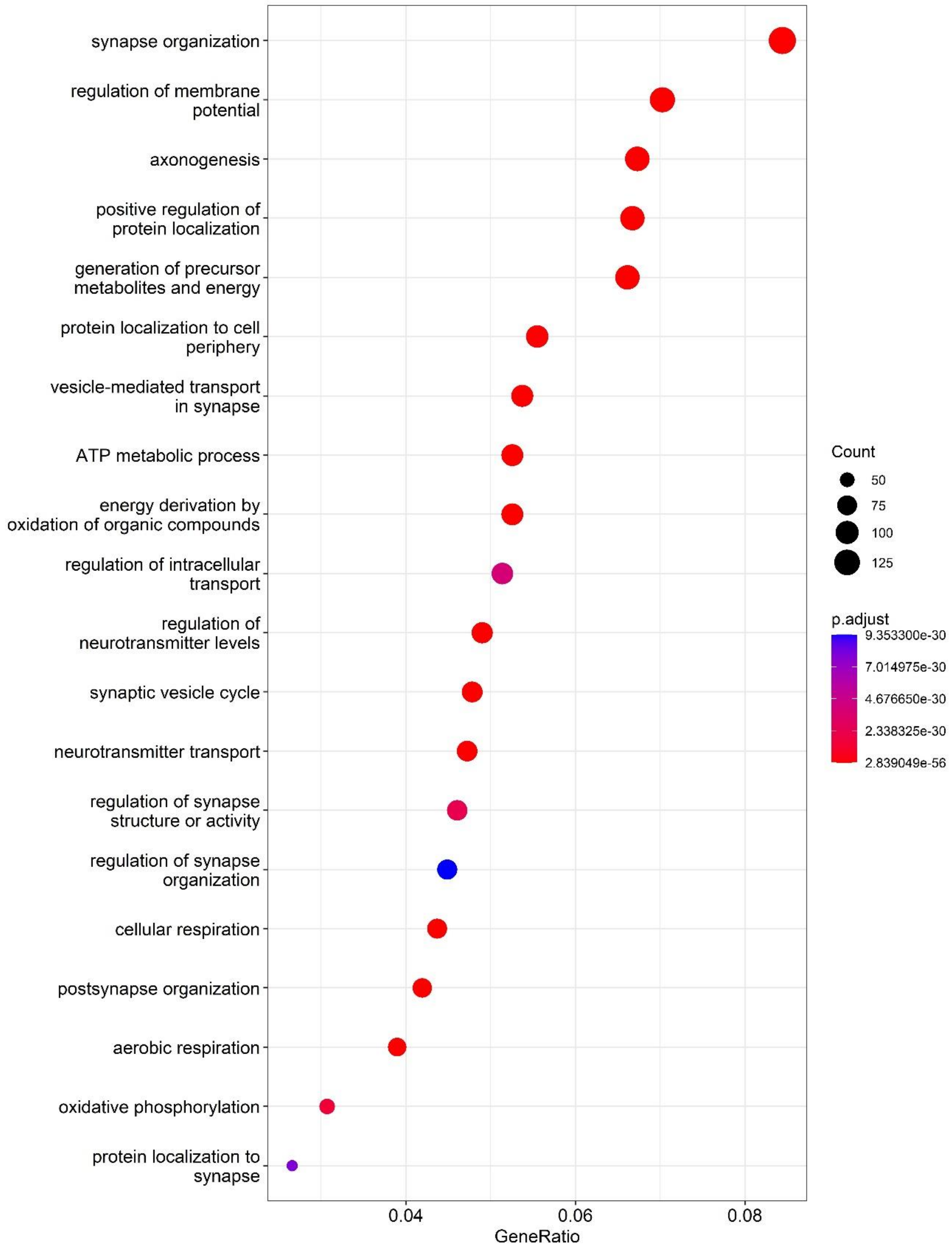


Supplementary Figure 13. Enriched gene ontology terms on 10X Visium mouse brain Posterior 1 data.

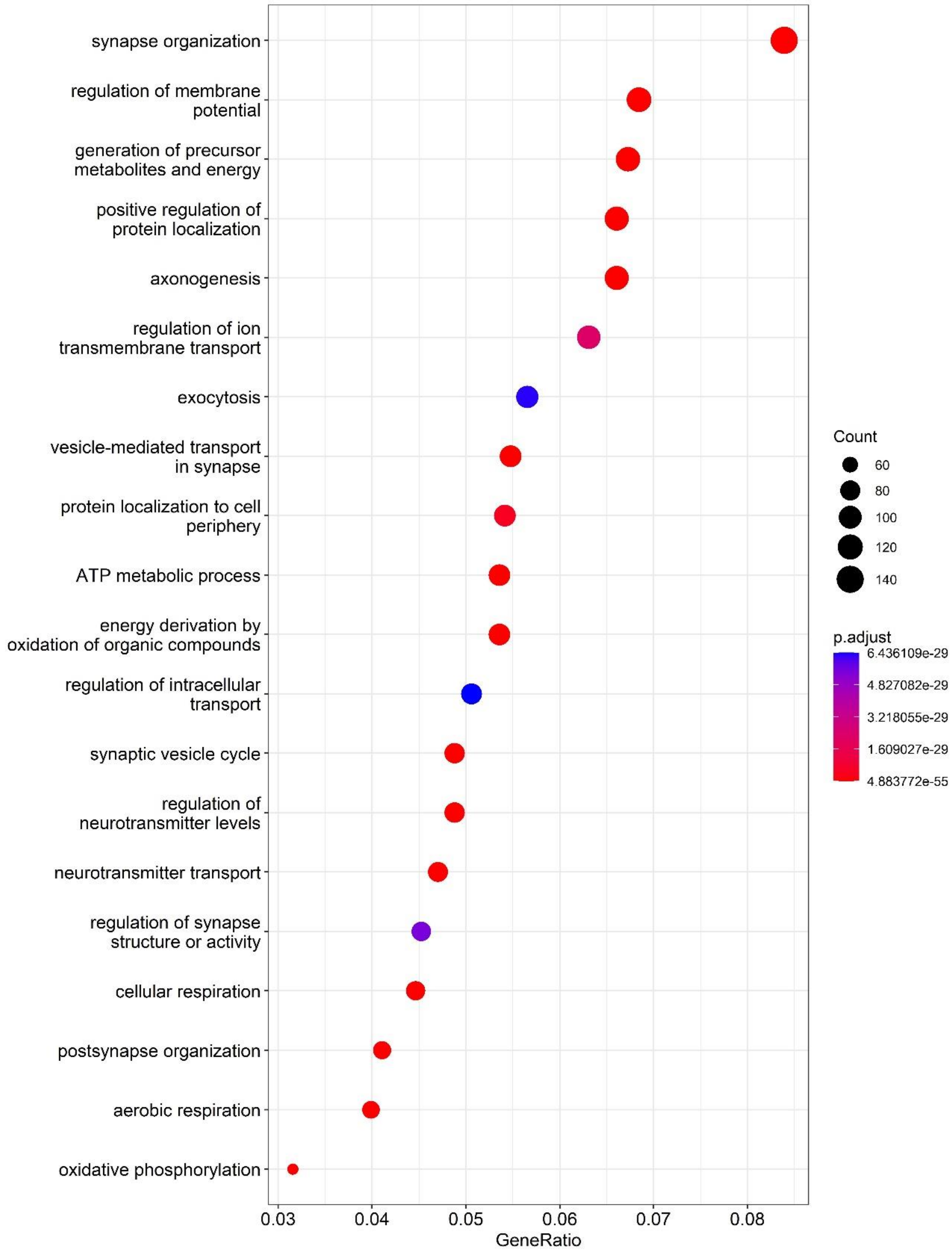


Supplementary Figure 14. Enriched gene ontology terms on 10X Visium mouse brain Posterior 2 data.

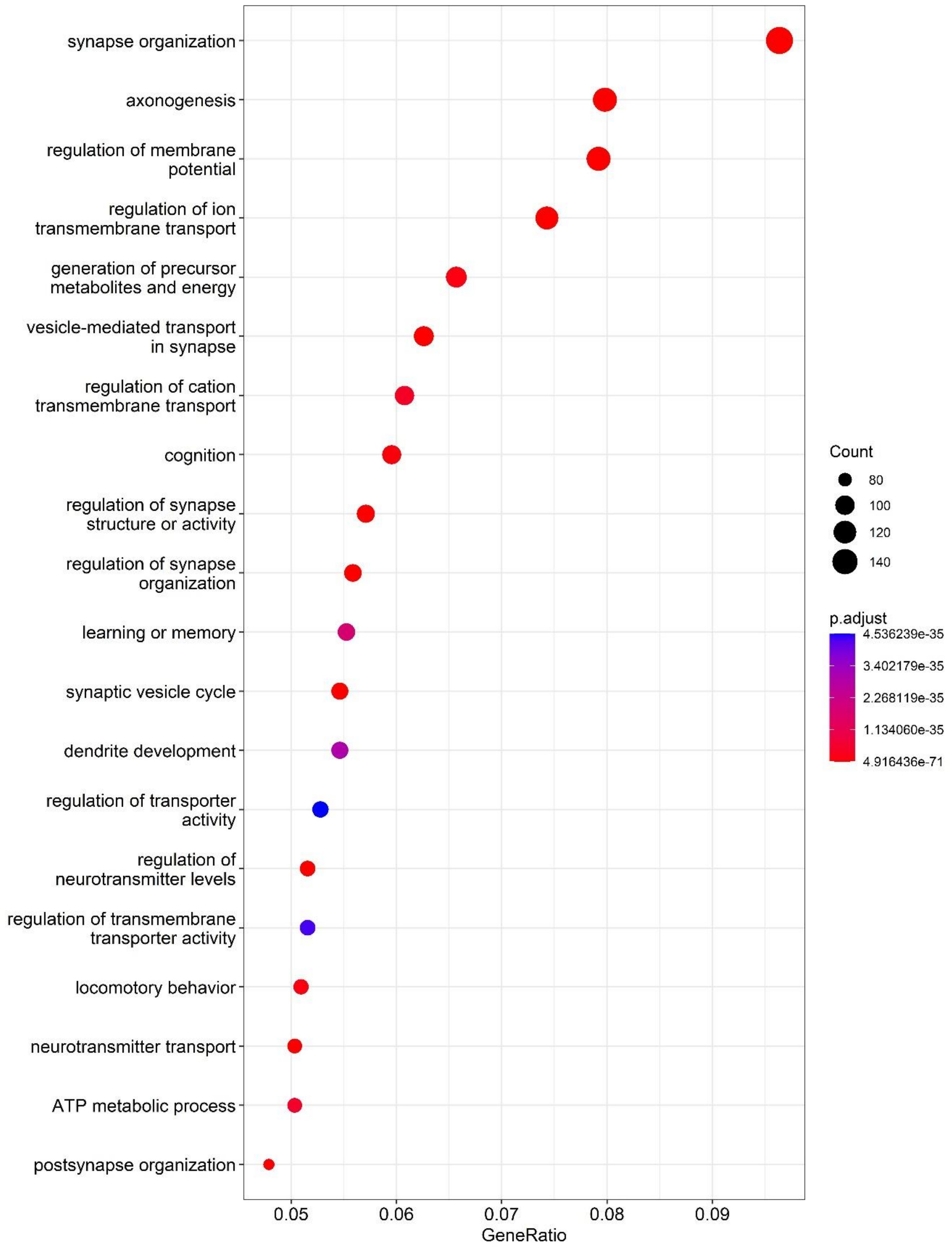


Supplementary Figure 15. Enriched gene ontology terms on Stereo-seq mouse brain data.
