## Supplementary Table 1 for "scBSP: A fast and accurate tool for identifying spatially variable genes from spatial transcriptomic data"

Small Sample

| FileName | N_Genes | N_Spots | scBSP_time | scBSP_mem | SPARKX_tim | SPARKX_me | SOMDE_tim | SOMDE_me | BSP_time | BSP_mem | nnSVG_time | nnSVG_mer | MoranI_tim | MoranI_me | SPARK_time | SPARK_men | spatialDE_ti | spatialDE_mem |
|---|---|---|---|---|---|---|---|---|---|---|---|---|---|---|---|---|---|---|
| 10000_3000.mtx | 10000 | 3000 | 2.96 | 390.3 | 25.266 | 174.3 | 381.6293 | 317.0326 | 44.14873 | 1148.29 | | | | | | | 162.633 | 1442.849 |
| 15000_3000.mtx | 15000 | 3000 | 4.415 | 583.8 | 37.9 | 258.4 | 574.0399 | 475.4176 | 61.64429 | 1721.135 | | | | | | | 165.2062 | 1672.325 |
| 2000_3000.mtx | 2000 | 3000 | 0.567 | 104.5 | 5.026 | 48.1 | 76.64874 | 63.59283 | 15.92668 | 231.7348 | 2163.284 | 552.2 | | | 4397.076 | 1096.6 | 164.9363 | 1076.614 |
| 20000_3000.mtx | 20000 | 3000 | 5.996 | 778 | 51.203 | 253 | 761.474 | 633.8351 | 79.52763 | 2293.996 | | | | | | | 166.9274 | 1900.842 |
| 25000_3000.mtx | 25000 | 3000 | 7.446 | 971.5 | 64.77 | 316 | 954.4873 | 792.1651 | 97.51806 | 2866.826 | | | | | | | 167.1937 | 2129.955 |
| 30000_3000.mtx | 30000 | 3000 | 8.837 | 1165.2 | 77.209 | 379.2 | 1151.572 | 950.6106 | 115.9553 | 3439.688 | | | | | | | 165.8316 | 2358.834 |
| 35000_3000.mtx | 35000 | 3000 | 10.358 | 1359.7 | 90.764 | 442.3 | 1335.703 | 1108.964 | 152.875 | 4012.531 | | | | | | | 163.859 | 2587.83 |
| 4000_3000.mtx | 4000 | 3000 | 1.265 | 157.6 | 10.077 | 72.4 | 153.0971 | 127.0834 | 22.7194 | 460.8731 | | | | | | | 161.8173 | 1168.053 |
| 40000_3000.mtx | 40000 | 3000 | 11.915 | 1553.3 | 103.285 | 505.4 | 1528.48 | 1267.357 | 153.4059 | 4585.376 | | | | | | | 165.6631 | 2816.826 |
| 6000_3000.mtx | 6000 | 3000 | 1.8 | 235.2 | 15.072 | 106.4 | 229.4393 | 190.4362 | 29.90774 | 690.0085 | | | | | | | 160.6732 | 1259.652 |
| 8000_3000.mtx | 8000 | 3000 | 2.339 | 312.6 | 20.26 | 140.4 | 313.408 | 253.9892 | 37.11095 | 919.1493 | | | | | | | 164.2563 | 1351.25 |
| 20000_1000.mtx | 20000 | 1000 | 2.337 | 268.2 | 49.819 | 119.1 | 701.5751 | 230.0325 | 28.28695 | 766.3082 | | | | | | | 38.40573 | 458.3429 |
| 20000_10000.mtx | 20000 | 10000 | 25.353 | 2581 | 56.027 | 840.3 | 666.8643 | 2047.486 | 365.9395 | 7640.971 | | | | | | | 2912.97 | 13811.45 |
| 20000_2000.mtx | 20000 | 2000 | 3.883 | 522.6 | 50.762 | 231 | 706.243 | 432.6319 | 52.72012 | 1530.147 | | | | | | | 75.25478 | 1053.819 |
| 20000_3000.mtx | 20000 | 3000 | 6.001 | 778 | 50.984 | 253 | 759.8372 | 633.8346 | 80.07932 | 2293.995 | | | | | | | 160.4913 | 1900.841 |
| 20000_4000.mtx | 20000 | 4000 | 7.945 | 1033.8 | 51.94 | 336.9 | 813.1299 | 836.4126 | 110.5932 | 3057.846 | | | | | | | 307.3235 | 2961.489 |
| 20000_500.mtx | 20000 | 500 | 1.15 | 140.1 | 49.696 | 63.1 | 738.8955 | 129.229 | 16.26396 | 384.3914 | 10204.38 | 111.5 | | | 5949.133 | 791.3 | 33.55481 | 229.4391 |
| 20000_5000.mtx | 20000 | 5000 | 13.123 | 1296 | 52.344 | 420.8 | 800.6574 | 1035.389 | 149.0541 | 3821.703 | | | | | | | 515.8349 | 4235.757 |
| 20000_6000.mtx | 20000 | 6000 | 15.469 | 1553.1 | 53.199 | 504.7 | 776.2947 | 1239.612 | 172.1554 | 4585.562 | | | | | | | 784.6533 | 5723.765 |
| 20000_7000.mtx | 20000 | 7000 | 13.648 | 1801.5 | 53.667 | 588.6 | 750.4188 | 1439.46 | 239.0506 | 5349.407 | | | | | | | 1126.834 | 7425.164 |
| 20000_8000.mtx | 20000 | 8000 | 15.547 | 2057.4 | 54.676 | 672.5 | 701.2221 | 1645.585 | 261.9124 | 6113.548 | | | | | | | 1655.442 | 9340.42 |
| 20000_9000.mtx | 20000 | 9000 | 17.447 | 2313.4 | 55.093 | 756.5 | 671.4571 | 1846.405 | 321.4031 | 6877.115 | | | | | | | 2278.284 | 11469.06 |

Large Sample

| FileName | N_Genes | N_Spots | Sparsity | scBSP_time | scBSP_mem | SPARKX_tim | SPARKX_me | SOMDE_tim | SOMDE_me | BSP_time | BSP_mem |
|---|---|---|---|---|---|---|---|---|---|---|---|
| 10000_100000_0.00 | 10000 | 100000 | 0.000503 | 2.969 | 380.2 | 27.234 | 75 | 1393.463 | 8029.981 | 7908.721 | 38239.99 |
| 15000_100000_0.00 | 15000 | 100000 | 0.0005 | 3.494 | 383.7 | 39.476 | 75.5 | 1933.177 | 12041.33 | 9194.3 | 57314.11 |
| 2000_100000_0.000 | 2000 | 100000 | 0.000792 | 2.236 | 374.6 | 6.74 | 76.2 | 447.0225 | 2718.714 | 7745.105 | 7721.379 |
| 20000_100000_0.00 | 20000 | 100000 | 0.000499 | 4.386 | 451.8 | 52.937 | 86.8 | 2589.442 | 16052.84 | 9501.628 | 76388.53 |
| 25000_100000_0.00 | 25000 | 100000 | 0.0005 | 4.758 | 539.1 | 65.687 | 92.5 | 3176.595 | 20064.31 | 8167.982 | 95462.37 |
| 30000_100000_0.00 | 30000 | 100000 | 0.0005 | 5.292 | 626.4 | 78.105 | 100.2 | 3763.907 | 24075.88 | 9697.285 | 114536.5 |
| 35000_100000_0.00 | 35000 | 100000 | 0.0005 | 5.816 | 714.1 | 90.685 | 122.2 | 4363.136 | 28087.36 | 8620.047 | 133610.6 |
| 4000_100000_0.000 | 4000 | 100000 | 0.000578 | 2.432 | 375.1 | 11.741 | 70 | 685.2663 | 3216.431 | 8884.178 | 15351.03 |
| 40000_100000_0.00 | 40000 | 100000 | 0.0005 | 6.175 | 801.5 | 102.96 | 122.3 | 4986.612 | 32098.88 | 10457.58 | 152684.8 |
| 6000_100000_0.000 | 6000 | 100000 | 0.000526 | 2.538 | 376.6 | 16.833 | 78.2 | 920.8802 | 4820.882 | 8939.694 | 22980.68 |
| 8000_100000_0.000 | 8000 | 100000 | 0.00051 | 2.747 | 377.3 | 21.806 | 75.1 | 1164.35 | 6425.42 | 7848.151 | 30610.33 |
| 20000_100000_0.00 | 20000 | 100000 | 0.000204 | 2.864 | 377.4 | 52.019 | 78 | 2614.055 | 16046.19 | 9418.226 | 76315.01 |
| 20000_100000_0.00 | 20000 | 100000 | 0.000301 | 3.283 | 379.2 | 51.942 | 83.7 | 2596.237 | 16048.38 | 8770.218 | 76388.26 |
| 20000_100000_0.00 | 20000 | 100000 | 0.0004 | 3.633 | 385.3 | 51.653 | 81.7 | 2593.898 | 16050.76 | 8276.581 | 76388.26 |
| 20000_100000_0.00 | 20000 | 100000 | 0.000499 | 4.037 | 451.9 | 52.632 | 91.5 | 2590.996 | 16052.84 | 9494.645 | 76388.25 |
| 20000_100000_0.00 | 20000 | 100000 | 0.0006 | 4.532 | 518.3 | 52.191 | 88.5 | 2589.64 | 16055.08 | 10125.71 | 76388.26 |
| 20000_100000_0.00 | 20000 | 100000 | 0.0007 | 5.097 | 584.5 | 52.545 | 102.7 | 2587.059 | 16057.31 | 9847.703 | 76388.53 |
| 20000_100000_0.00 | 20000 | 100000 | 0.000799 | 5.23 | 650.4 | 52.247 | 98.3 | 2588.556 | 16059.53 | 9519.689 | 76388.26 |
| 20000_100000_0.00 | 20000 | 100000 | 0.000899 | 5.8 | 716.3 | 51.875 | 95.5 | 2582.33 | 16061.92 | 10215.08 | 76388.26 |
| 20000_100000_0.00 | 20000 | 100000 | 0.000999 | 5.977 | 782.3 | 52.013 | 98.2 | 2588.031 | 16064.2 | 8793.59 | 76388.26 |
| 20000_100000_0.00 | 20000 | 99997 | 0.0005 | 4.293 | 452.3 | 52.294 | 90.7 | 2478.264 | 16052.55 | 8062.604 | 76385.97 |
| 20000_150000_0.00 | 20000 | 149994 | 0.0005 | 6.424 | 671.9 | 52.186 | 127.7 | 4430.081 | 24071.5 | 22029.59 | 114577 |
| 20000_200000_0.00 | 20000 | 199993 | 0.0005 | 7.464 | 775 | 52.533 | 153.2 | 7007.333 | 32076.45 | 37992.73 | 152769 |
| 20000_250000_0.00 | 20000 | 249988 | 0.0005 | 11.793 | 1112 | 52.717 | 199.6 | 10535.55 | 40098.73 | | |
| 20000_300000_0.00 | 20000 | 299987 | 0.0005 | 14.491 | 1331.9 | 53.603 | 221.1 | 15148 | 48128.72 | | |
| 20000_350000_0.00 | 20000 | 349983 | 0.0005 | 17.574 | 1551.9 | 53.737 | 260.4 | 21284.56 | 56153.99 | | |
| 20000_400000_0.00 | 20000 | 399977 | 0.0005 | 20.482 | 1772 | 53.318 | 268.9 | 27474.18 | 64166.57 | | |
| 20000_50000_0.000 | 20000 | 49997 | 0.0005 | 2.728 | 232.5 | 50.169 | 50.2 | 1231.547 | 8032.782 | 2094.328 | 38192.99 |
