## Supplementary Table 2 for "scBSP: A fast and accurate tool for identifying spatially variable genes from spatial transcriptomic data"

| | | | | | | | | | |
|---|---|---|---|---|---|---|---|---|---|
| 1 | Dataset | Sample | Original_Ge | Permuted_ | Cell_Count | scBSP_time | scBSP_men | SPARK_tim | SPARK_mem |
| 2 | Xenium_Ca | Sample1_R | 313 | 10016 | 166363 | 158.886 | 19375.1 | 74.983 | 7797.7 |
| 3 | Xenium_Ca | Sample1_R | 313 | 10016 | 118691 | 113.107 | 14035 | 57.362 | 5679.4 |
| 4 | Xenium_Ca | Sample2 | 313 | 10016 | 118691 | 113.2 | 14034 | 57.637 | 5677.3 |
| 5 | Xenium_M | Xenium_V1 | 248 | 10168 | 162033 | 269.236 | 22765 | 112.061 | 12392.1 |
| 6 | Xenium_M | Xenium_V1 | 248 | 10168 | 154654 | 253.801 | 21602 | 105.674 | 11589.4 |
| 7 | Xenium_M | Xenium_V1 | 248 | 10168 | 158047 | 276.299 | 22049.1 | 106.353 | 11766.6 |
| 8 | CosMx_Lur | Lung5_Rep | 980 | 10780 | 100292 | 67.911 | 12206.6 | 40.474 | 3227.6 |
| 9 | CosMx_Lur | Lung5_Rep | 960 | 10560 | 106660 | 78.695 | 13351.8 | 45.71 | 3773.9 |
| 10 | CosMx_Lur | Lung5_Rep | 960 | 10560 | 100264 | 64.139 | 11552.7 | 38.246 | 2868.5 |
| 11 | CosMx_Lur | Lung6 | 960 | 10560 | 93795 | 57.694 | 10587.4 | 39.183 | 3082.8 |
| 12 | CosMx_Lur | Lung9_Rep | 960 | 10560 | 91972 | 57.741 | 10880.1 | 37.894 | 2872.7 |
| 13 | CosMx_Lur | Lung9_Rep | 960 | 10560 | 150504 | 155.879 | 17198.6 | 45.527 | 3741.7 |
| 14 | CosMx_Lur | Lung12 | 960 | 10560 | 73997 | 50.492 | 9356.4 | 34.944 | 2421.9 |
| 15 | CosMx_Lur | Lung13 | 960 | 10560 | 82843 | 54.977 | 9930.7 | 38.09 | 2745.5 |
| 16 | Steroseq_M | Mouse_bra | 26177 | | 38746 | 25.112 | 7063.1 | 47.19 | 1192.5 |
| 17 | Steroseq_M | Mouse_olfa | 26145 | | 107416 | 53.525 | 14851.1 | 51.144 | 1442.9 |
| 18 | Steroseq_M | Mouse_olfa | 23815 | | 104931 | 65.375 | 14730.1 | 46.467 | 1377.8 |
| 19 | HDST | HDST | 19950 | | 181367 | 5.951 | 681.8 | 33.156 | 100.7 |
| 20 | Visium | anterior1 | 20467 | | 2696 | 6.33 | 1197.2 | 38.3 | 482.2 |
| 21 | Visium | anterior2 | 20368 | | 2825 | 5.78 | 1155.5 | 38.2 | 446.5 |
| 22 | Visium | posterior1 | 20524 | | 3353 | 4.85 | 1081.1 | 36.89 | 479.1 |
| 23 | Visium | posterior2 | 20328 | | 3293 | 4.78 | 1040.1 | 36.86 | 455.1 |
